# Gene flux drives functional changes in an ancient supergene in *Formica lemani*

**DOI:** 10.64898/2026.10.05.756876

**Authors:** Zul Alam, Giulia Scarparo, German Lagunas-Robles, Daniela Zarate, Jessica Purcell, Alan Brelsford

## Abstract

Pinpointing the genetic elements underlying a phenotype within a supergene can be difficult because hundreds of linked genes can show similar associations with phenotype. Rare recombination between supergene haplotypes can help identify those genetic elements by changing haplotype-phenotype combinations. Here we investigate the genetic underpinnings of colony queen number and body size in the ant *Formica lemani*. Throughout the *Formica* genus, single-queen colonies typically carry M haplotypes, while multi-queen colonies also carry P haplotypes. In *F. lemani*, we found two P haplotypes homologous to those described in the closely related *F. cinerea*, but unlike in *F. cinerea*, where P_1_ is restricted to multi-queen colonies, *F. lemani* P_1_ is common in both single- and multi-queen colonies. This deviation suggests that P_1_ may carry novel SNP combinations allowing single queen colony compatibility. By comparing haplotypes within *F. lemani* and with *Formica cinerea*, we searched for candidate monogyny variants resulting from gene flux from the M onto the P_1_ haplotype, or with P_1_-specific mutations. We found 82 genes associated with candidate variants, of which two are among the fourteen previously suggested candidates in other species, suggesting that several different genes may underlie queen number across the genus. We also identified a supergene associated with smaller queens and males, which is shared with *F. cinerea* and in interchromosomal linkage disequilibrium with the queen-number supergene. Together, these results provide a rare example of how an ancient supergene can change in function, and a step toward identifying the genes within a supergene responsible for its functions.

## Introduction

Identifying the functional genetic elements underlying a phenotype can be difficult due to the presence of linked sites which show similar associations with phenotype. This difficulty is further amplified in regions of low recombination, such as inversions, which can include hundreds of linked genes. When these regions of low recombination contain multiple functional genetic elements and control discrete complex polymorphic phenotypes they are called “supergenes” (Thompson & Jiggins, 2014). Supergenes are known to underlie traits such as mimetic coloration in butterflies (e.g., Joron *et al*., 2011; Kunte *et al*., 2014; Nishikawa *et al*., 2015), flower morph in angiosperms (e.g., Li *et al*., 2016; Gutiérrez-Valencia *et al*., 2022), and colony queen number in ants (e.g., Wang *et al*., 2013; Purcell *et al*., 2014). Although inversions, and other structural variants, greatly suppress recombination, rare recombination events still occur. Recombination can occur in inversion heterozygotes via gene conversion or double crossovers, processes collectively referred to as gene flux (Navarro *et al*., 1997; Guerrero *et al*., 2012).

Gene flux has two consequences that can help identify candidate genes for supergene functions. First, gene flux gradually erodes neutral variation between haplotypes while selection preserves differentiation at sites contributing to maintaining the alternate phenotypes (Navarro *et al*., 1997; Guerrero *et al*., 2012). For example, the wing coloration supergene in the butterfly *Heliconius numata* shows evidence of recombination between haplotypes, but high differentiation at the gene *cortex (Jay et al., 2021)*. The second, less-studied consequence of gene flux is its ability to move candidate genes from one haplotype onto another, creating new haplotype-phenotype combinations. In the butterfly *Danaus chrysippus*, recombination moved an allele for dark coloration, *yellow*, onto the alternate haplotype, helping distinguish this candidate gene from the many linked variants within the supergene (De-Kayne *et al*., 2025). Thus, gene flux can reveal candidates both through the differentiated regions that resist homogenization and through phenotype-associated variants that cross haplotype backgrounds.

The ant genus *Formica*, with its ancient supergene underlying queen number, provides an opportunity to use both consequences of gene flux to uncover candidate genes (Brelsford *et al*., 2020; Purcell *et al*., 2021). Throughout the *Formica genus*, individuals from monogyne (single-queen) colonies consistently carry only the “M” haplotype and individuals from polygyne (multi-queen) colonies carry the “P” haplotype, although the within-colony distribution of haplotypes may vary (Purcell *et al*., 2014; Lagunas-Robles *et al*., 2021, 2025; McGuire *et al*., 2022; Pierce *et al*., 2022; Scarparo *et al*., 2024). In comparisons of distantly related species, gene flux has homogenized most of this ancient supergene, leaving small clusters of SNPs which consistently differentiate the M and P haplotypes (Brelsford *et al*., 2020; Purcell *et al*., 2021). Most conserved SNPs fall within the gene *knockout* (Brelsford *et al*., 2020), a transcription factor required for motor neuron development (Hartmann *et al*., 1997). There are also a few species with apparent genotype-phenotype mismatches such as *F. podzolica*, where some polygyne colonies have the monogyne *knockout* allele (Purcell *et al*., 2025). Similarly, in *F. paralugubris* the P haplotype is not found in polygyne colonies (Lagunas-Robles *et al*., 2025), but P-specific duplications in the highly polygynous *F. paralugubris* and *F. aquilonia* have made their way onto the M (Sigeman *et al*., 2025a). The ancient origin of the supergene coupled with the diversity in genotype-phenotype associations allow for comparative studies that can shed light on the evolutionary history of the supergene (Brelsford *et al*., 2020; Purcell *et al*., 2021).

Brelsford *et al*. (2020) found *F. cinerea* and *F. lemani* to each have two P haplotypes. Building on this, Scarparo *et al*. (2023) confirmed the presence of two P haplotypes (P_1_ and P_2_) on chromosome 3, both associated with polygyny, and discovered a second supergene on chromosome 9 underlying queen and male miniaturization. They further highlighted strong linkage disequilibrium (LD) between the two supergenes, such that nearly all individuals carrying the multi-queen P_2_ haplotype on chromosome 3 also carry the miniaturizing 9r haplotype on chromosome 9 (Scarparo *et al*., 2023). The presence of two P haplotypes in *F. lemani* suggests it may be a useful comparison for identifying candidate genes and determining whether the chromosome 9 supergene is found in other species.

In this study, we aimed to examine the genetic underpinnings of colony social structure and queen, male and worker size in *Formica lemani*. We began by investigating the genotype distribution in monogyne and polygyne colonies. Surprisingly, we found a P haplotype (P_1_) to be common in monogyne colonies. Following this we searched for candidate genes involved in monogyny, comparing the P and M haplotypes in *F. lemani* and *F. cinerea* to look for gene flux events and *de novo* mutations which may have caused the *F. lemani* P_1_ haplotype to become compatible with mongyny. Next, we searched for a miniaturizing supergene on chromosome 9 and assessed its linkage disequilibrium to the social supergene on chromosome 3. Finally, we determined whether the miniaturizing supergene haplotype on chromosome 9 (and, separately, the associated P_2_ haplotype on chromosome 3) evolved independently in *F. cinerea* and *F. lemani*.

## Methods

### Field Sampling

The black ant *Formica lemani* is widely distributed throughout the mountain ranges of Europe and Asia, and forms both monogyne (single-queen) and polygyne (multi-queen) colonies (Seppä *et al*., 2009; Seifert, 2018). We sampled *F. lemani* from a single population in Bosco Gurin, Switzerland (46.316° N, 8.492° E) between 2018 and 2021. We collected at least 10 workers per colony, and if observed, we sampled up to 10 males and 10 gynes (virgin queens), collectively known as alates (winged individuals which reproduce). Ants were only sampled from directly under rocks that cover colony entrances. Samples were stored in 100% ethanol. Sample metadata can be found in Table S1.

### DNA Extraction

We extracted DNA using two different protocols. For samples collected during 2018, we used a modified version of the Qiagen DNeasy Blood & Tissue Kit protocol for insect tissue (McGuire *et al*., 2022). For each colony, we extracted DNA from the whole body of up to 6 workers and the heads of up to 8 gynes and males when available. Colonies which were sampled over multiple years had several samples extracted per year. Each sample was ground with sterile pestles after immersion in liquid nitrogen. The pulverized tissues were then digested overnight in a mix of Buffer ATL and Proteinase K. Deviations from the Qiagen protocol included completing the second wash with 70% ethanol in place of AW2, using alternatively sourced spin columns (BPI-tech.com), and eluting in 30 µL of Buffer AE.

For samples collected after 2018, we ground the ant tissues as described above, and after overnight digestion in Buffer ATL and proteinase K, we transferred the supernatant to the QIAcube HT/QIAxtractor robot for washing and elution following the QiaAmp 96 DNA automated protocol. We eluted the DNA in 100 µL of EB Buffer.

### RadSeq Library Preparation

For the *F. lemani* samples sequenced using double-digest restriction site associated DNA sequencing (RADseq), we followed the methods from Brelsford *et al*., (2016). First, we digested 6 ul of DNA per sample using MseI and PstI restriction enzymes. We then ligated a universal MseI adaptor and uniquely coded PstI adaptors to each sample. Next, we removed small fragments using Serapure magnetic beads (Rohland & Reich, 2012) or Omega magnetic beads (OMEGA Bio-Tek) in a 0.8:1 (beads: sample) ratio. We amplified the DNA in four separate 5 ul PCR reactions (with Q5 Hot Start DNA Polymerase) and added plate-specific indexed Illumina primers. We then pooled the replicate PCR products for each sample and ran a final PCR cycle. Each PCR product was run on a 1.5% agarose gel, and samples successfully amplified were pooled together. Finally, we removed small fragments using the same beads and ratio as before. Samples were sequenced by the University of California San Diego’s Institute for Genomic Medicine on an Illumina NovaSeq 6000 (S4 flow cell) or by Novogene Inc. on an Illumina Hiseq X, with 150bp paired end reads.

### Seqwell Library Preparation

We also prepared 44 *F. lemani* individuals for whole genome resequencing using the Seqwell plexWell LP-384 library kit following manufacturer’s protocol. Samples were sequenced by Novogene Inc on a Illumina NovaSeq 6000 with 150bp paired end reads.

### Public Data

We used the 33 *F. cinerea* individuals from Lagunas-Robles *et al*. (2026) (BioProject PRJNA1425027), and two *F. cinerea* individuals and one *F. truncorum* male from Brelsford *et al*. (2020) (BioProject PRJNA557080)

### Bioinformatics

#### Data preparation

We used Stacks v2.1 and Stacks v2.60 to demultiplex samples from 2018 and 2019-2021, respectively (Catchen *et al*., 2011). We used PEAR v0.9.6 to merge overlapping paired-end reads and remove adaptor sequences (Zhang *et al*., 2014), then aligned reads to the *F. selysi* reference genome (Brelsford *et al*., 2020; NCBI GCA_009859135.1) using BWA-mem2 2.2.1 (Vasimuddin *et al*., 2019). We called SNPs using Samtools mpileup 1.14 (Li, 2011).

For the RadSeq dataset, we filtered genotypes for biallelic variants (--max-alleles 2), a minimum depth of 8 (--minDP 8), loci with under 10% of missing data (--max-missing 0.9), sites with a minor allele count of at least 2 (--mac 2), and removed indels (--remove-indels) using VCFtools 0.1.17 (Danecek *et al*., 2011). Duplicated regions were removed by calculating the proportion of heterozygosity at each locus (--hardy) in haploid males, then excluding sites from all individuals that appeared heterozygous in at least 5% of males as described in Scarparo *et al*. (2023). Our final dataset counted 1043 individuals and 23,189 sites.

For the whole genome dataset, we filtered genotypes for a minimum depth of 2 (--minDP 2), loci with under 50% missing data (--max-missing 0.5), sites with a minor allele count of at least 2 (--mac 2), and removed indels (--remove-indels) using VCFtools 0.1.17 (Danecek *et al*., 2011). Our final dataset counted 79 individuals and 5,942,763 sites.

#### Linking colony social form to supergene genotype on chromosome 3

To investigate the association between social form and chromosome 3 genotype, we used the Radseq data to assign genotypes based on a principal component analysis (PCA) using the 1,279 loci on the chromosome 3 VCF with PLINK v1.90b6.25 (Purcell *et al*., 2007). We also considered F_IS_ values for each individual (--het) on chromosome 3 using VCFtools, where individuals with negative values are considered heterozygous while positive values are considered homozygous. Although males are typically haploid, we sampled some diploid males and assigned them diploid genotypes. We then extracted genotypes with bcftools v1.19 (bcftools query) (Danecek *et al*., 2021) and compared the frequency of reference and alternative alleles relative to the *F. selysi* reference genome built from a pool of M males, to make a preliminary assessment about whether haplotypes were M-like or P-like. Next, we looked at the genetic differentiation (F_ST_) between each haplotype pair by selecting only homozygous individuals and estimating Weir and Cockerham’s F_ST_ between each pair of genotypes using VCFtools. We visualized the F_ST_ comparison using the R package qqman (Turner, 2018).

We used the Wang estimator in CoAncestry 1.0.2.0 (Wang, 2011) to compute pairwise intracolony relatedness and infer colony social form on the Radseq samples. To allow an assessment of the social form independent from the supergenes, we excluded markers on chromosomes 3 and 9. We also removed all the males and excluded colonies with fewer than five diploid individuals (workers and gynes). Our final, refiltered dataset for CoAncestry resulted in 795 individuals, 104 colonies and 23,071 SNPs. Distributions of intracolony relatedness were visualized using the R packages ggridges (Wilke, 2026) and ggplot2 (Wickham, 2016).

Following Scarparo *et al*. (2023), we considered colonies with all pairwise estimates ≥ 0.6 as monogyne monandrous; colonies with a bimodal distribution and at least 40% of pairwise relationships ≥ 0.6 and none <0.2 as monogyne polyandrous; colonies with at least one pairwise relationship ≤ 0.1 were considered polygyne.

Finally, we tested the association between genotype and colony social form by fitting a generalized linear model with a binomial error distribution and logit link function. Since we did not collect alates from every colony, we only included worker genomes in this analysis, and we fit the model at the colony level with one row per colony (n = 102 colonies). We used the social form of the colony encoded as a binomial (0 = monogyne, 1 = polygyne) as the response variable . We included two fixed effects in the same model, P_1_ presence and P_2_ presence, which we encoded as binomials (0 = absence in the colony, 1 = presence), allowing us to assess the association of each haplotype with social form while accounting for the other. We ran the model using the “glm” function in base R v4.5.2 (R Core Team, 2025). We tested the association between our fixed effects and social form using a type III Anova with the “Anova” function from the car package (Fox & Weisberg, 2019).

#### SNPs underlying social structure on chromosome 3

Unlike in *F. cinerea*, *F. lemani’s* P_1_ haplotype is common in monogyne colonies (Fig. 2), despite topology weighting suggesting homology between the P_1_ haplotypes (Fig. S3; Supplemental Methods). Given the P haplotype’s history of gene flux (Brelsford *et al*., 2020), it is possible that such events moved “M-associated” SNPs to *F. lemani*’s P_1_ haplotype, leading to the observed genotype distribution. To search for these SNPs (hereafter “candidate monogyny variants”, or “CMoVs”) resulting from gene flux, we used our whole-genome dataset. We first looked for SNPs which had a fixed difference (F_ST_ > 0.975) between other P haplotypes (*F. lemani* P_2_, and *F. cinerea* P_1_ and P_2_) and M haplotypes (from *F. lemani* and *F. cinerea*). This list was subsetted for sites where at least 90% of the *F. lemani* individuals homozygous for P_1_ have the same allele as the M haplotype. SNPs with > 70% missing data across individuals were excluded. We also included four sites where at least 90% of P_1_P_1_ individuals are heterozygous, representing likely P_1_ specific duplications where one copy is fixed for the same allele as the M haplotype. We retained a total of 208 CMoVs of putative gene flux origin.

As the observed distribution of P_1_ in *F. lemani* may also be due to *de novo* mutations, we also considered *F. lemani* P_1_-specific nonsynonymous *de novo* mutations as CMoVs. Using our whole genome dataset, we selected SNPs with F_ST_ > 0.975 when comparing *F. lemani* P_1_P_1_ individuals to *F. cinerea* of all genotypes and *F. lemani* individuals without the P_1_. We then determined the functional impact (i.e. synonymous, non-synonymous) of the SNP using SNPeff v5.2e (Cingolani *et al*., 2012) and only retained nonsynonymous mutations for a total of 9 *de novo* CMoVs.

#### Gene annotation

For the 208 gene flux CMoVs and 9 *de novo* mutation CMoVs, we assessed the position of the SNP relative to the reference genome annotation from the global ant genome alliance (Vizueta *et al*., 2025), and categorized the SNP as either intergenic or intragenic. For intragenic SNPs, we recorded the gene the SNP fell within and used SNPeff (Cingolani *et al*., 2012) to annotate and predict the SNPs’ functional effect. For intergenic SNPs, we recorded the nearest upstream and downstream genes and calculated the distance from the SNP to the gene. We determined gene ID and function using BLASTX using the -db flag to restrict our database to *D. melanogaster*. We only assigned a *D. melanogaster* ortholog for E values lower than 1×10^-5^, and excluded others from downstream analyses.

We then sorted the candidate genes into five tiers reflecting the strength of the evidence linking them to colony queen number. Tier one genes have been proposed as candidates for colony social organization in a previous *Formica* study and also carry a CMoV. Tier two genes carry a gene flux CMoV that SNPeff predicts to cause a missense mutation. Tier three genes carry a *de novo* CMoV predicted to cause a missense mutation. Tier four genes carry a gene flux CMoV predicted to be intronic or synonymous. Tier five genes are the nearest upstream or downstream gene to an intergenic CMoV. Genes meeting the criteria for more than one tier were assigned to the highest tier that applied.

#### GO term analysis

To examine Gene Ontologies and search for overrepresented GO terms associated with our genes we used the PANTHER Overrepresentation Test (Mi *et al*., 2019; Thomas *et al*., 2022) with the Biological Processes Complete annotation dataset. Enriched GO categories were identified using the Fisher’s Exact Test with an FDR correction. Only genes with an assigned ortholog were considered. We tested for functional class overrepresentation in genes overlapping or adjacent to the 208 gene flux CMoVs, and separately tested for functional class overrepresentation in genes containing the 9 *de novo* mutation CMoVs. We additionally tested for functional class overrepresentation among all genes in the supergene region (Scaffold03: 2,000,000-12,500,000).

#### Searching for a supergene on chromosome 9

We assessed the presence of a supergene on chromosome 9 by performing a PCA using the 1,192 SNPs on chromosome 9, and assigned genotypes to individuals based on clustering on the PCA, F_IS_ values, and F_ST_ as described for chromosome 3.

We noticed that individuals with the P_2_ haplotype on chromosome 3 almost always had the same haplotype of chromosome 9. We provisionally called this haplotype of chromosome 9 “9r” and the other haplotype “9a”following the naming convention in *F. cinerea* (Scarparo *et al*., 2023). We confirmed the linkage disequilibrium between haplotypes in *F. lemani* using a Fisher’s exact test using 9r haplotype presence/absence and P_2_ presence/absence with the fisher.test function in base R (R Core Team, 2025).

#### Chromosome 9 Homology

To investigate the phylogenetic relationships between haplotypes of the supergene on chromosome 9 and confirm the names we assigned to each haplotype, we conducted topology weighting using twisst2 (Martin, 2026). We created a new vcf consisting of all loci on chromosome 9 from all homozygotes for chromosome 9 haplotypes from the whole-genome *F. lemani* (n=34) and *F. cinerea* (n=35) datasets and included one *F. truncorum* male (Brelsford *et al*., 2020, SRS10116540) for polarization. Since we only included homozygotes and haploids, we called pseudo-haplotypes by setting ploidy to 1 when calling SNPs. We polarized our data using sticcs (Martin, 2026), and filtered the vcf for minimum depth of 1 (--minDP 1), loci with under 50% of missing data (--max-missing 0.5), sites with a minor allele count of at least 2 (-- mac 2), and retained bi-allelic loci (--max-alleles 2) and removed indels (--remove-indels) using VCFtools 0.1.17 (Danecek *et al*., 2011). We then inferred genealogies and counted subtree topologies using twisst2 (Martin, 2026), and calculated topology weights and visualized using the plot_twisst.R (Martin, 2026) script in R.

We additionally identified SNPs associated with the 9r haplotype across species. Homologous supergenes should be enriched for 9r specific sites which are shared across species. We identified sites at which an allele was present at a frequency greater than 0.95 in both the *F. lemani* 9r and *F. cinerea* 9r, then removed sites with that allele present at a frequency greater than 0.05 in either 9a group. We retained 950 shared 9r associated SNPs genome wide. Of these SNPs, 925 are found on chromosome 9 between 2.39-9.45 Mb and the 25 remaining SNPs are found on chromosome 3.

#### Linking body size to supergene genotype on chromosome 9

To study the association between body size and supergene haplotype, we measured the head width of 64 gynes and queens, 107 males, and 160 workers using a Leica DMC2900 camera mounted on a Leica S8APO at 25× magnification. Head width is a good proxy for body size as it has a strong positive correlation with several body segment dimensions in *Formica* species (Tawdros *et al*., 2020). All individuals measured for head size were sequenced via Radseq.

We tested for an association between body size and chromosome 9 genotype by fitting three independent linear mixed models (LMM), one for each caste (gynes/queens, males and workers) in R (R Core Team, 2025). We used head width as the response variable, chromosome 9 genotypes as a fixed effect and colony as a random effect. We determined pairwise P values using emmeans v2.0.0 with a Tukey post-hoc (Lenth & Piaskowski, 2025). We also fit three LMMs, one for each caste, to test for an association between body size and chromosome 3 genotype.

#### M_2_ haplotype analysis

To determine whether the *F. lemani* M_2_ haplotype is homologous to the *F. cinerea* M_D_ haplotype, we performed F_ST_, windowed PCA, topology weighting, and linkage mapping (Supplementary methods; Fig. S6).

## Results

### The queen number supergene on chromosome 3

#### Chromosome 3 harbors four supergene haplotypes

We determined there are four supergene haplotypes on chromosome 3 via genomic clustering (PCA) and heterozygosity (via F_IS_) analyses. Six distinct clusters were distinguished on Principal components (PC) 1 and 2 suggesting the presence of three different haplotypes (Fig. 1a). However, PC3 showed the presence of ten distinct clusters, revealing a fourth haplotype (Fig. 1b). We determined there are four homozygous and six heterozygous clusters via our heterozygosity analysis (F_IS_; Fig. S1). We found all 10 possible genotype combinations in workers, gynes and queens, and all four haplotypes in haploid males. Assessment of reference and alternate allele frequencies relative to the *F. selysi* M reference genome suggested that two haplotypes were “M-like” (M_1_ and M_2_) and two were “P-like” (P_1_ and P_2_) (Fig. S2).

**Figure 1:**
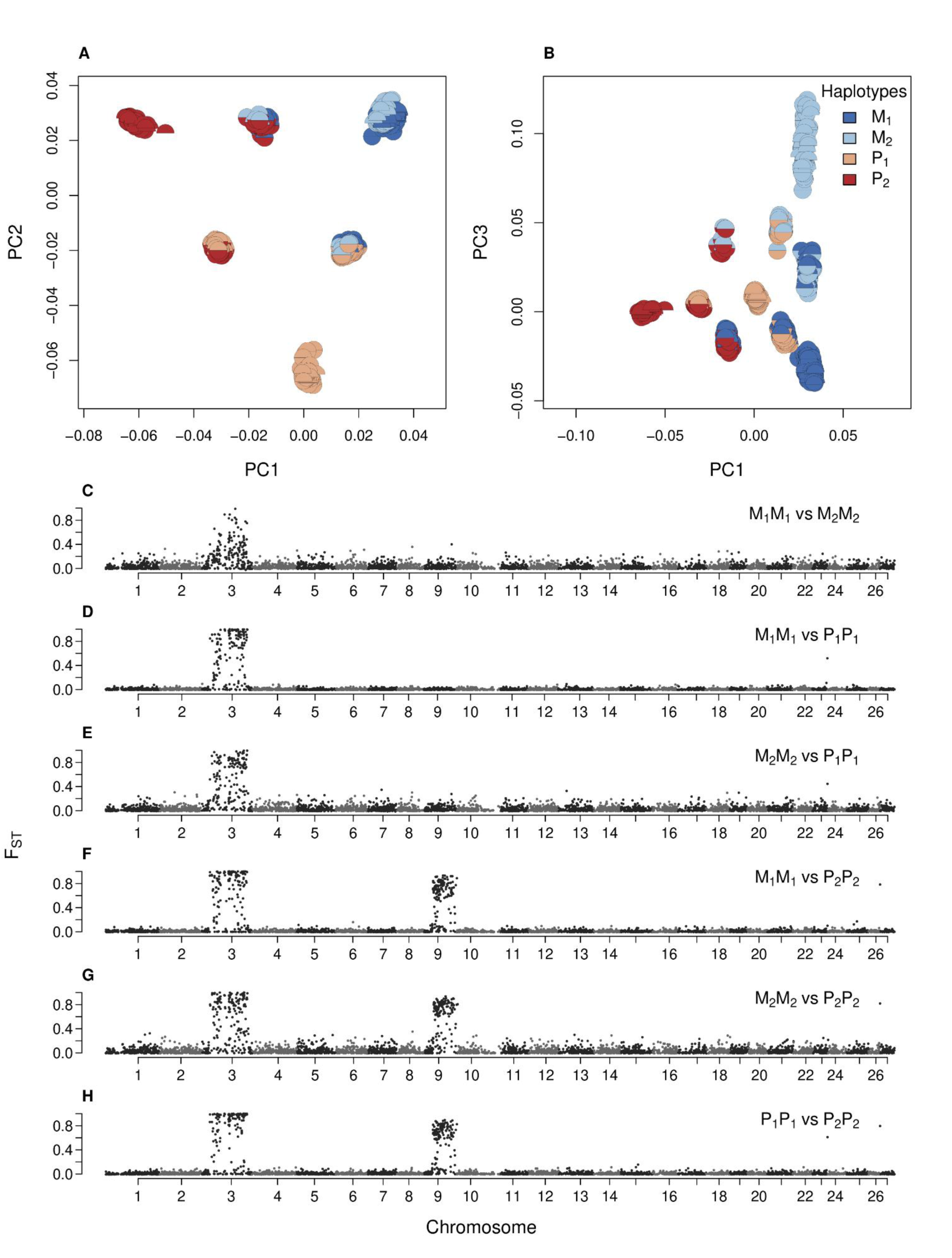
PCA and F_ST_ reveal four haplotypes on chromosome 3 in *F. lemani* (A) PC1 (17.07% variance explained) and PC2 (12.23%) calculated with RADseq markers separate six clusters, consistent with three haplotypes. (B) PC3 (5.07%) reveals a fourth haplotype. Each half circle is one chromosome 3 haplotype. Diploid females and males appear as two halves of a full circle and haploid males as a single half. (C) M_1_ and M_2_ are differentiated across the entire supergene region. (D and E) The M and P_1_ haplotypes are differentiated across most of chromosome 3. (F-H) The P_2_ haplotype on chromosome 3 is differentiated from all other haplotypes across most of chromosome 3 and chromosome 9. F_ST_ was calculated in 10kb windows with RADseq markers. Alt Text: Two scatter plots showing genomic clustering on chromosome 3. Below are six genome scans showing fixed differences between haplotypes.

The analysis of F_ST_ between haplotypes revealed high differentiation between the two M haplotypes (M_1_ and M_2_), and high differentiation between the P_1_ and both M haplotypes spanning most of chromosome 3 from 2 Mbp to 12.5 Mbp (Figure 1c). The P_2_ shows high differentiation both between 2 Mbp and 12.5 Mbp on chromosome 3 and on an 7 Mbp region on chromosome 9 when compared with the other haplotypes. Using twisst2 (Martin, 2026) we computed topology weightings for the *F. lemani* and *F. cinerea* P haplotypes. We found that within the supergene region on chromosome 3, the most common topology groups both P_1_ haplotypes together and both P_2_ haplotypes together rather than grouping by species, while the species tree was most common outside the supergene (Fig. S3).

#### Loose association of the ‘social’ supergene with colony queen number

To determine if the supergene on chromosome 3 underlies queen number in *F. lemani* we assigned social structure to colonies based on intracolony pairwise relatedness. Of the 104 analyzed colonies, 31 colonies are monogyne monandrous, 13 are monogyne polyandrous (multiple patrilines), and 60 are polygyne (Fig. 2a).

**Figure 2:**
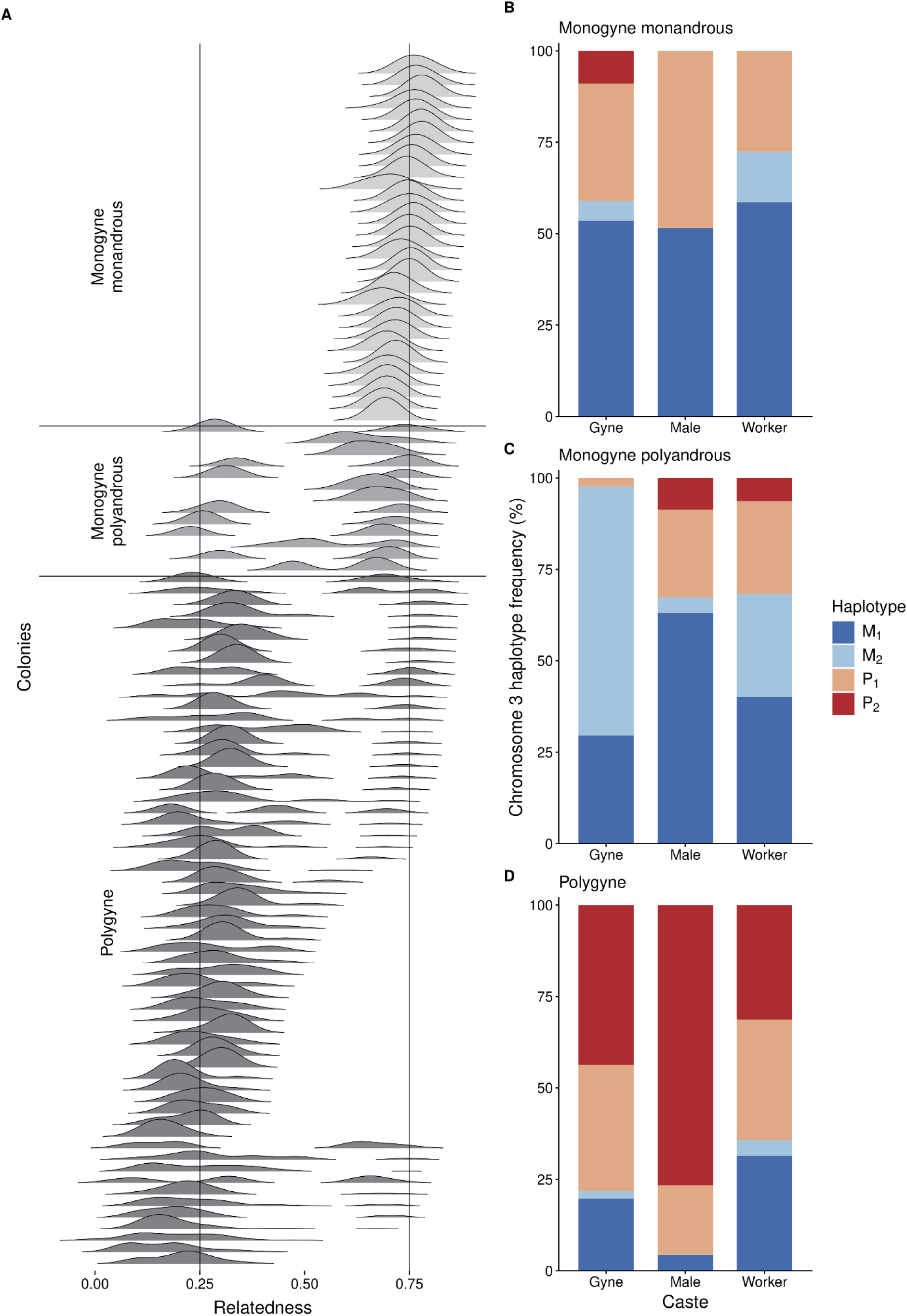
Haplotype distribution across social forms shows that P_2_ is concentrated in polygyne colonies while P_1_ is common in both monogyne and polygyne colonies. (A) Ridgeline plot showing pairwise relatedness values for workers in 104 colonies sequenced via RADseq. 31 colonies are monogyne monandrous, 13 are monogyne polyandrous (multi-male), and 60 are polygyne. (B-D) Haplotype frequency for each caste by social form sequenced via RADseq. (B) Monogyne monandrous colonies have mostly M_1_ and P_1_ individuals. (C) Monogyne polyandrous colonies have mostly M_1_, M_2_ and P_1_ individuals. (D) Polygyne colonies have mostly P_2_ and P_1_ individuals. Alt Text: Left shows a ridgeline plot displaying pairwise intra-colony relatedness values as a distribution for each colony. The right shows three stacked bar plots displaying haplotype frequencies in monogyne monandrous, monogyne polyandrous, and polygyne colonies.

We found P_2_ haplotype presence to be strongly associated with polygyny (χ^2^_1_= 46.83, P = 7.727e-12). Out of 60 polygyne colonies, 43 had at least one worker with the P_2_ haplotype (Fig. S4) whereas only two of the 44 monogyne colonies (both inferred to be polyandrous) harbored individuals with the P_2_ haplotype (Fig. 2a, 2b). The P_2_ haplotype is also the most common haplotype carried by gynes and males from polygyne colonies (Fig. 2d).

Accounting for P*_2_* presence, we also found P_1_ presence to be weakly associated with polygyny (χ^2^ = 6.45, P = 0.01113). While the P_1_ haplotype is common in polygyne colonies (58/60 polygyne colonies harbor the P_1_ haplotype) it is also common in monogyne colonies (33/44 monogyne colonies harbor the P_1_ haplotype) (Fig. S4). Based on worker genotypes, we infer that 17 monogyne, monandrous colonies are headed by a M_1_/P_1_ heterozygous queen, mated with an M_1_ male (n=13), P_1_ male (n=3), or M_2_ male (n=1; Fig. S4). We note that several of these colonies produced sexual offspring, indicating that they were mature.

#### Candidate genes for colony queen number

The prevalence of the P_1_ haplotype in *F. lemani* monogyne colonies provides an opportunity to investigate the genes underlying variation in colony queen number. The functional difference in the *F. lemani* P_1_ compared to the other P haplotypes could result from gene flux of monogyne-associated alleles from M onto P_1_, or from *de novo* mutations that disrupt genes responsible for polygyne social structure on the P_1_. We found 208 candidate monogyny variants (CMoVs) resulting from gene flux from *F. lemani’s* Ms to P_1_. We identified an additional nine nonsynonymous *de novo* mutations on *F. lemani’s* P_1_ (all missense) which we considered as CMoVs. All CMoVs resulting from gene flux, including synonymous mutations and intergenic SNPs, are considered because they originated on a monogyne-associated M haplotype. Of the *de novo* mutations specific to *F. lemani’s* P_1_, only the nine nonsynonymous mutations will be considered CMoVs because the majority of *de novo* intergenic and synonymous SNPs are expected to be neutral (Eyre-Walker & Keightley, 2007).

The 208 CMoVs resulting from gene flux are distributed throughout the supergene region on chromosome 3; 110 are intragenic and 98 intergenic (Fig. 3; Table S2). These SNPs occur in 105 10kb bins, possibly representing 105 different gene conversion or double recombination events between P_1_ and M in *F. lemani*. We also identified nine CMoVs resulting from missense *de novo* mutations on *F. lemani* P_1_.

**Figure 3:**
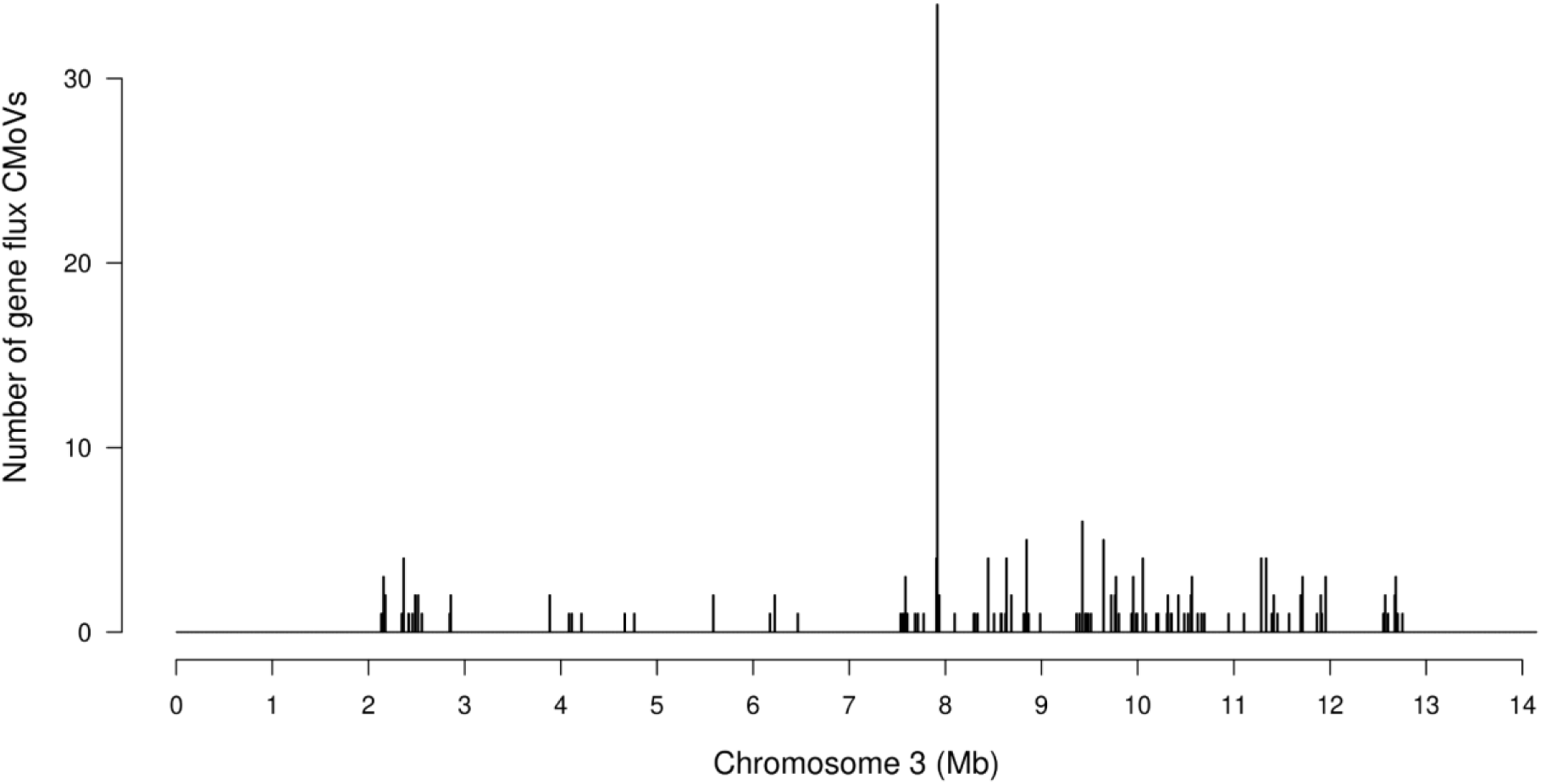
Candidate Monogyny Variants (CMoVs) resulting from gene flux from the *F. lemani* M haplotype onto the P_1_ haplotype are distributed throughout the supergene region on chromosome 3. These 208 CMoVs, identified via whole genome resequencing data, occupy 105 10-kbp windows. Alt Text: Bar plot showing the number of gene flux CMoVs distributed across chromosome 3

CMoVs are associated with 82 genes, and we separated those genes into five tiers (Table S3). Tier one genes (n=2) have been previously suggested as candidate genes, and have an associated CMoV. *Knockout*, a storkhead-box transcription factor with a role in neuron development (Hartmann *et al*., 1997), carries two intronic gene-flux CMoVs. *G9a*, which is involved in gene expression, dendrite morphogenesis, larval locomotion, and memory (Kramer *et al*., 2011), carries a *de novo* missense CMoV.

Tier two genes (n=5) carry a gene-flux CMoV that causes a missense mutation (Table S3). These include *sev*, which alone accounts for four of the eight within-gene missense CMoVs and is required for photoreceptor specification (Basler & Hafen, 1988).

Tier three genes (n=6) carry a nonsynonymous *de novo* CMoV (Table S3). Among these, *LamC*, involved in muscle tissue morphogenesis (Dialynas *et al*., 2010) carries three *de novo* missense CMoVs.

Tier four comprises 23 genes, 20 of which have *D. melanogaster* orthologs, with an intronic or synonymous gene-flux CMoV (91 CMoVs). Several genes are involved in anatomical structure development (n = 9) and nervous system development (n = 6). The 46 Tier five genes (31 of which have *D. melanogaster* orthologs) are the nearest upstream and downstream gene to an intergenic CMoV. Several of these genes are also involved in anatomical structure development (n = 13) and nervous system development (n = 7).

We found genes associated with a gene flux CMoV to be enriched for 12 different GO Biological Process terms. Eight of those terms were also enriched in the supergene region as a whole including anatomical structure development, and nervous system development (Table S4). Four GO terms were uniquely enriched in genes associated with a gene flux CMoV. These include regulation of dendrite development, muscle organ development, regulation of cell projection organization, and regulation of plasma membrane bounded cell projection organization. Genes with a *de novo* CMoV (n = 4 with *D. melanogaster orthologs*) were not enriched for any GO terms.

#### Chromosome 9 has a queen miniaturizing supergene

The elevated F_ST_ on chromosome 9 when comparing the P_2_ haplotype to other haplotypes on chromosome 3 suggested that *F .lemani* may harbor a second supergene on chromosome 9, as described in its close relative *F. cinerea* (Scarparo *et al.,* 2023). The PCA performed with markers on chromosome 9 revealed three distinct clusters: two with homozygous individuals and one with heterozygous individuals, corresponding to two alternative haplotypes (9a and 9r) (Fig. 4a).

**Figure 4:**
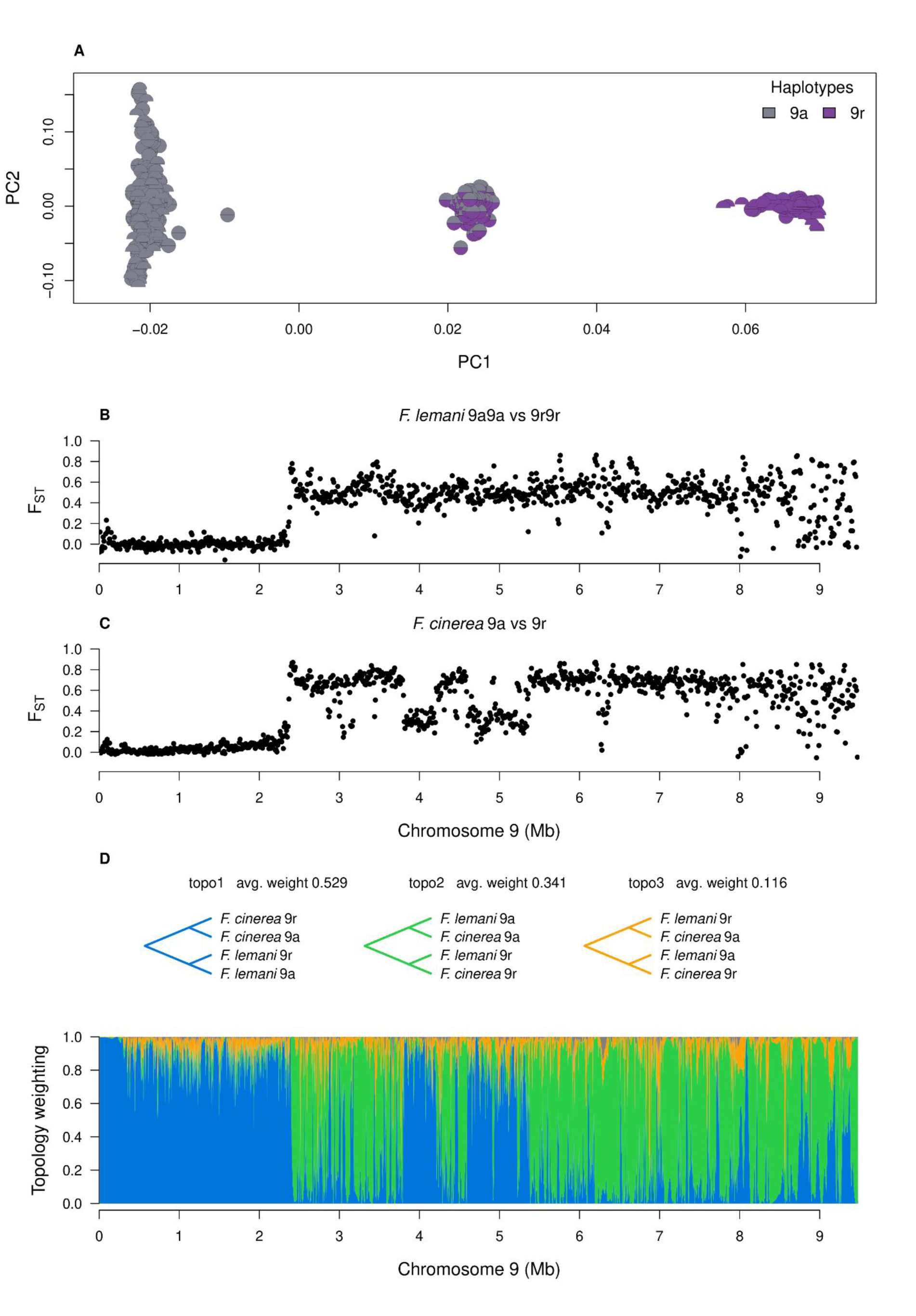
Chromosome 9 has two haplotypes in *F. lemani*, which are homologous to two *F. cinerea* haplotypes. A) PCA of RADseq SNPs shows two haplotypes on chromosome 9 in *F. lemani*. PC1 explains 23.79% of total variance and PC2 explains 1.27%. (B-C) F_ST_ between 9a and 9r haplotypes shows the same elevated region from ∼2.4 Mbp to 9.4 Mbp in both *F. lemani* and *F. cinerea*. F_ST_ was calculated from WGS data in 10 kbp windows. (D) Topology weightings calculated from whole genome resequenced data across chromosome 9 for *F. lemani* and *F. cinerea* 9a and 9r. The most common topology from ∼2.4 Mbp to 9.4 Mbp groups by haplotypes rather than species Alt Text: Top panel shows a scatter plot showing genomic clustering on chromosome 9. Below that are two genome scans showing differentiation between the 9a and 9r haplotypes. At the bottom is a genome scan showing topology weights on chromosome 9.

We determined that *F. lemani’s* 9r is homologous to *F. cinerea’s*. The genomic supergene region spans about 7 Mbp, from about 2.4 Mbp to 9.4 Mbp in both species (Fig. 4). Using twisst2 (Martin, 2026) we computed topology weightings for the four haplotypes (Fig. 4d). We found the species tree to be the most common topology in the recombining region of chromosome 9, while within the supergene region we found the tree grouping haplotypes together to be the most common topology (Fig. 4d). The regions within the supergene where the species tree is most common are likely sites of double crossover or gene conversion.

Additionally, we searched for 9r haplotype-specific SNPs which are shared across species, and found the 9r haplotypes share 925 SNPs in the supergene region but none elsewhere on chromosome 9, further supporting homology between the haplotypes (Fig. S5). We additionally found 25 within the chromosome 3 social supergene, likely due to the linkage disequilibrium between 9r and P_2_, and none elsewhere in the genome.

The P_2_ haplotype on chromosome 3 and the 9r haplotype on chromosome 9 are in high linkage disequilibrium (Fisher’s exact test: OR = 162.74, P < 2.2e-16; levels: presence-absence P_2_; presence-absence 9r). Scarparo *et al*. (2023) called the presence of 9r without P_2_ (or vice versa) a mismatch (e.g. P_2_X-9a9a or XX-9a9r, where X can be either M or P_1_). Following that definition, at population-level, we found 6.8% of our samples (71/1043) are mismatched (Table S5), all being workers (11% of workers, 71/640). However, our full dataset includes individuals from colonies whose queens carry neither P_2_ nor 9r, and therefore cannot show mismatch combinations. This means our initial estimate of mismatch frequency is likely an underestimate. We therefore restricted our dataset to colonies with at least one P_2_ or 9r haplotype (because cotransmission cannot occur without at least one of the haplotypes present). Under this criterion, 23.5% (71/301) of workers are mismatched in *F. lemani*. We reanalyzed the genotypes from Scarparo *et al*. (2023) using the new criteria and found 11.8% (19/161) of *F. cinerea* workers are mismatched (Table S5). Among the 219 males and 155 gynes we genotyped (92 and 74 from a colony with at least one haplotype respectively), no mismatches were observed.

We found that chromosome 9 genotype was significantly associated with gyne size (F_2_ = 14.09, P = 0.000167) and male size (F_2_ = 22.09, P = 2.531e-06), but not worker size (F_2_ = 2.76, P = 0.0669). 9r9r gynes are 14.5% smaller (t_26.4_ = 4.80, P = 0.0002), and 9a9r gynes are 9.2% smaller (t_18.4_ = 3.44, P = 0.0077) than 9a9a gynes, respectively. 9a males are 9.2% larger than 9r males (t_29.1_ = 5.79, P < 0.0001). 9r9r workers are 7.4% smaller (t_139.4_ = 2.12, P = 0.0901) and 9a9r workers are 3.4% smaller (t_93.2_ = 1.60, P = 0.2515) than 9a9a workers, but worker size differences are not statistically significant.

We also tested the association between chromosome 3 genotypes and body size, and found a significant association with queen size (F_7_ = 3.71, P = 0.005946), male size (F_6_ = 7.85, P = 1.258e-05), and worker size (F_9_ = 2.82, P = 0.005536). Genotypes with P_2_ were often smaller than those without (e.g., P_1_P_2_ gynes vs M_1_M_1_ t_24.9_ = 3.91, P = 0.0123; see Table S6 for full list).

## Discussion

Most of the previously studied *Formica* species show two alternative haplotypes at the social supergene (Purcell *et al*., 2014; Brelsford *et al*., 2020; Pierce *et al*., 2022; McGuire *et al*., 2022; Scarparo *et al*., 2024). Expanding upon Brelsford *et al*. (2020)’s initial observation, we found four distinct haplotypes (M_1_, M_2_, P_1_, P_2_) on chromosome 3 associated with queen number in *F. lemani*. These findings resemble the recently documented supergene diversity in the closely related species *F. cinerea* (Scarparo *et al*., 2023), although our investigation also uncovered notable distinctions. Unlike in all other *Formica* species investigated (Purcell *et al*., 2014; Brelsford *et al*., 2020; Lagunas-Robles *et al*., 2021; McGuire *et al*., 2022; Pierce *et al*., 2022; Scarparo *et al*., 2023, 2024) we found the P_1_ haplotype to be common in monogyne colonies, possibly due to recent gene flux from the M to the *F. lemani* P_1_ haplotype. By comparing the *F. lemani* P_1_ sequence to all other chromosome 3 haplotypes across *F. lemani* and *F. cinerea* we identified novel candidate genes for queen number. We also found a second supergene on chromosome 9 that has two haplotypes, the 9a associated with larger queen and male sizes and the 9r associated with small queens and males. Finally, we found that LD between the two supergenes on chromosomes 3 and 9 is strong in *F. lemani,* though not as strong as the LD described in *F. cinerea* (Scarparo *et al*., 2023). *F. lemani* shows that ancient supergene systems can continue to evolve in two ways, by taking on novel phenotype associations by changing the contents of haplotypes, and by expanding to incorporate new traits.

### Gene flux reveals candidate genes

In most *Formica* species, the M and P haplotypes are consistently associated with monogyny and polygyny, respectively (Purcell *et al*., 2014; Brelsford *et al*., 2020; McGuire *et al*., 2022; Pierce *et al*., 2022; Scarparo *et al*., 2023). Here, we found that the M_1_, M_2_, and P_2_ haplotypes in *F. lemani* have generally maintained their strong association with colony social form. In contrast, the P_1_, while weakly associated with polygyny, is also present in 75% of monogyne colonies (Table 1). Given the extent of P_1_ presence in *F. lemani’s* monogyne colonies, we suggest that a sequence change to the P_1_ may allow it to exist in monogyne colonies. Two non-mutually-exclusive proximate mechanisms could have allowed P_1_ to become compatible with monogyny: gene flux copying monogyny-associated alleles from an M haplotype onto P_1_, or P_1_-specific *de novo* mutations that disrupt loci required for polygyny.

**Table 1.**

| Species where P is found in monogyne colonies | Proportion of monogyne colonies with at least one individual with a P haplotype | Citation |
| --- | --- | --- |
| *F. neoclara* | 10% (1/10) | McGuire *et al.* 2022 |
| *F. cinerea* | 8.3% (5/60) | Scarparo *et al.* 2023 |
| *F. aserva* | 4.8% (1/21) | Scarparo *et al.* 2024 |
| *F. podzolica* | 15% (13/86) | Purcell *et al.* 2025 |
| *F. lemani* | 75% (33/44) | This study |

Gene flux is often discussed in the context of erosion in supergenes (Navarro *et al*., 1997; Guerrero *et al*., 2012; Brelsford *et al*., 2020; Berdan *et al*., 2021; Charlesworth, 2023; Meyer *et al*., 2024). Rare recombination events between haplotypes gradually homogenizes the haplotypes while selection preserves differentiation at functionally important loci (Navarro *et al*., 1997; Guerrero *et al*., 2012). Across the *Formica* genus, this slow process has left the M and P haplotypes strongly differentiated within each species, but only consistently differentiated over a small fraction of the supergene between species (Brelsford *et al*., 2020; Purcell *et al*., 2021). In this study we considered the role of gene flux from a different perspective. Rather than asking which loci resisted gene flux, we identified sites at which P_1_ carries the M-associated allele while the other P haplotypes, which are strongly associated with polygyny, retain the alternative allele. If some of these alleles affect social form, gene flux may have modified the function of P_1_ rather than only eroding divergence.

Comparable gene flux events have occurred in a few other supergene systems. In the African Monarch butterfly, *Danaus chrysippus,* a gene flux event moving the *yellow* allele from the “orientis” haplotype to a “klugii” background allowed researchers to confirm *yellow’s* involvement in wing coloration (De-Kayne *et al*., 2025). In the Atlantic cod, *Gadus morhua*, a gene flux event moved a *vitellogenin* allele cluster, likely affecting egg buoyancy, from the stationary ecotype to the migratory ecotype background (Matschiner *et al*., 2022). Finally, *F. paralugubris* appears to have lost the P haplotype while retaining the polygyne phenotype (Lagunas-Robles *et al*., 2025; Sigeman *et al*., 2025a), likely as a result of gene flux of P-associated gene duplications onto an M background (Sigeman *et al*., 2025a). In contrast, the *F. lemani* P_1_ is common in monogyne colonies possibly because it acquired M associated alleles. These cases illustrate how exchange between haplotypes can generate new haplotype-phenotype combinations. We used this process in *F. lemani* to attempt to identify genes which may be responsible for P_1_’s change in phenotype.

We found 208 candidate monogyny variants (CMoVs) resulting from gene flux, and 9 P_1_-specific *de novo* mutation CMoVs, but it is unlikely that all of them affect social form. The queen number supergene already has a long history of rare recombination/gene conversion without changing the association of the P haplotype with polygyny (Brelsford *et al*., 2020). Many recombination events may simply be neutral or even beneficial, if they reduce mutational load on P_1_ with only a few affecting queen number. Additionally, many of the fixed differences between haplotypes are likely to be neutral as well (Kimura, 1968). We therefore expect only a small subset of genes to affect colony queen number, possibly a single gene or just a few acting together. We present a ranked list of candidate genes for colony queen number (Table S3).

Of the 14 previously suggested candidate genes for queen number in *Formica* (Brelsford *et al*., 2020; Purcell *et al*., 2021; Sigeman *et al*., 2025a; Boulain *et al*., 2026), we only found two to have any associated candidate monogyny variants (CMoVs): *Knockout* and *G9a*.

Brelsford *et al*. (2020) suggested *Knockout* as a candidate gene because it consistently resisted erosion due to gene flux across 15 species spanning the *Formica* genus. It is a storkhead-box transcription factor involved in neuron development in *Drosophila* (Hartmann *et al*., 1997). They also suggested *G9a*, which resisted erosion in 14 of 15 species (Brelsford *et al*., 2020). G9a is involved in gene expression, dendrite morphogenesis, larval locomotion, and memory (Kramer *et al*., 2011). We found *Knockout* to have two intronic gene flux CMoVs and two gene flux CMoVs (1.5kb away), and found G9a to have a missense *de novo* CMoV and five nearby gene flux CMoVs. Brelsford *et al*. (2020) identified candidate genes as those which resist gene flux, whereas we identified candidate genes as those which experienced gene flux in a haplotype with an altered phenotype. Discovering two candidate genes in parallel with both methods supports the likelihood that they contribute to the queen number phenotype.

While cross-species evidence suggests that these genes are involved in determining social structure, their associations may not be universal. For example, *Knockout* doesn’t appear to underlie social form in a few species. In *F. podzolica*, P haplotypes carrying the M-associated *Knockout* allele are still associated with polygyny (Purcell *et al*., 2025). Similarly, in the highly polygynous species *F. aquilonia* and *F. paralugubris*, the P-specific *Knockout* allele was absent (Sigeman *et al*., 2025a). These examples highlight the lineage-specific nature of the social supergene, and open the possibility for *F. lemani*-specific candidate genes. Candidate genes identified in one species should therefore not be assumed to apply to others without further evidence.

We also identified several strong candidates which have not been mentioned in previous studies (Table S3). Tier two genes (n = 5) have experienced gene flux of a missense mutation from M to P_1_, and are the strongest new candidates for monogyny. The three genes with known functions are involved in photoreceptor specification, transcription corepression, and acetyl-CoA metabolism (Basler & Hafen, 1988; Oba *et al*., 2005). Changes in sensory development could alter responses to social cues, while changes in transcriptional regulation, or metabolism could influence development. Tier three genes (n = 6) have *de novo* P_1_-specific mutations and are also strong candidates. The three genes with known functions are involved in muscle tissue morphogenesis and eye development (Delaney *et al*., 1991; Yoshida *et al*., 2005; Dialynas *et al*., 2010). Changes to sensory development may also affect responses to social cues while tissue morphogenesis can affect development. Genes with intronic or synonymous gene flux CMoVs (Tier four, n = 23) and those upstream or downstream of an intergenic CMoV (Tier five, n = 46) may also underlie monogyny through regulatory effects. Social supergene haplotypes are known to differ in gene expression and regulatory element activity in other ant species (Pierce *et al*., 2022; Jones *et al*., 2025; Boulain *et al*., 2026).

We found that genes associated with gene flux CMoVs have GO terms relating to development and neurogenesis, but these categories are enriched in the supergene as a whole. Unique to the candidate genes are GO terms related to dendrite and cell projection organization and muscle organ development. Interestingly, there is a lack of odorant-binding proteins (OBPs) or odorant receptors (ORs) associated with a CMoV. In our dataset, gFselCSP10 (chemosensory protein) and *pino* (involved in olfaction) are the only genes similar to OBPs/ORs associated with CMoVs (Rollmann *et al*., 2005). The fire ant *Solenopsis invicta* carries nine OBPs in its supergene, six of which have protein differences between haplotypes (Pracana *et al*., 2017).

The independently evolved supergene in the desert ant *Cataglyphis niger* shares three OBPs with *S. invicta* (Lajmi *et al*., 2026), including *GP-9*, the second described candidate gene in *S. invicta* (Ross & Keller, 1998; Keller & Ross, 1998). Although *Formica’s* social supergene lacks OBP or OR candidate genes or enrichment observed in *S. invicta or C. niger,* we found enrichment for genes related to brain development, which could influence how sensory cues are processed.

By comparing haplotypes between and within *F. cinerea* and *F. lemani*, we identified many recent gene conversion or double crossover events between P_1_ and M in *F. lemani*. The distribution of gene flux SNPs, with just a few per window in windows spread across the supergene, suggests that gene conversion (which affects small blocks of DNA) is more common than double crossover events (which affect larger blocks of DNA) (Navarro *et al*., 1997). These gene flux SNPs may also be due to recurrent mutations, but the gene flux rate is likely orders of magnitude faster than the mutation rate even in the inverted region (Navarro *et al*., 2000).

### A homologous queen size supergene

We determined that *F. lemani’s* 9r haplotype is homologous to that of *F. cinerea*. Both supergenes span the same region of chromosome 9 and the most common tree topology within this region groups individuals by haplotype rather than species. The 9r haplotypes also share 925 9r-specific SNPs within the supergene region, but none elsewhere on chromosome 9. *F. lemani* and *F. cinerea* last shared an ancestor over 13 million years ago (Borowiec *et al*., 2021), suggesting that the queen miniaturization supergene may be at least that old. This differs from independent recruitment of the same chromosome, which has occurred with the M_D_ haplotype between five Nearctic *Formica* species and *F. cinerea* (Lagunas-Robles *et al*., 2026) and between the social supergenes of *Solenopsis* and *Cataglyphis* (Lajmi *et al*., 2026). Introgression from one species to another could produce a similar pattern, such as in the social supergene of *Solenopsis* (Helleu *et al*., 2022) or the Y chromosome between the ninespine and Amur stickleback (Dixon *et al*., 2019). Several *Formica* species show evidence of hybridization and gene flow (Kulmuni *et al*., 2010; Seifert *et al*., 2010; Satokangas *et al*., 2023), including *F. cinerea* (Purcell *et al*., 2016). However, to our knowledge, no publications report hybridization or gene flow between *Formica* species as diverged as *Formica lemani* and *Formica cinerea* (∼13 MYA; Borowiec *et al*., 2021)

*F. lemani* gynes carrying at least one copy of 9r are 9.2% to 14.5% smaller than 9a9a homozygotes, while haploid 9r males are 8.4% smaller than 9a males. The 14.5% difference between 9r9r and 9a9a gynes aligns with the magnitude seen in other ant groups where the larger morph is called macrogyne and the smaller morph microgyne (Wolf & Seppä, 2016). The smaller 9.2% difference between 9a9r and 9a9a gynes still far exceeds the average ∼2% difference between monogyne and polygyne sexuals in *Formica* (Sundström, 1995; Rosset & Chapuisat, 2007; Meunier & Chapuisat, 2009).

While we have low statistical power to separate the effects of 9r and P_2_ due to the high degree of linkage disequilibrium between the haplotypes, there are several lines of evidence suggesting that 9r contributes to miniaturization more than P_2_. First, because Scarparo *et al*. (2023) demonstrated that 9r has a stronger direct effect on queen size than P_2_ via GWAS, it is most parsimonious that 9r rather than P_2_ miniaturizes queens and males in *F. lemani* as well. Second, 9r9r homozygous gynes are smaller than 9a9r heterozygotes, with an average head width of 1.48 ± 0.04mm; P_2_P_2_ homozygous gynes average 1.53 ± 0.04mm and are not significantly smaller than P_2_ heterozygotes. This pattern could be due to the few P_2_P_2_ gynes which are 9r heterozygotes, bringing up the average P_2_P_2_ size (Fig. 5).

**Figure 5.**
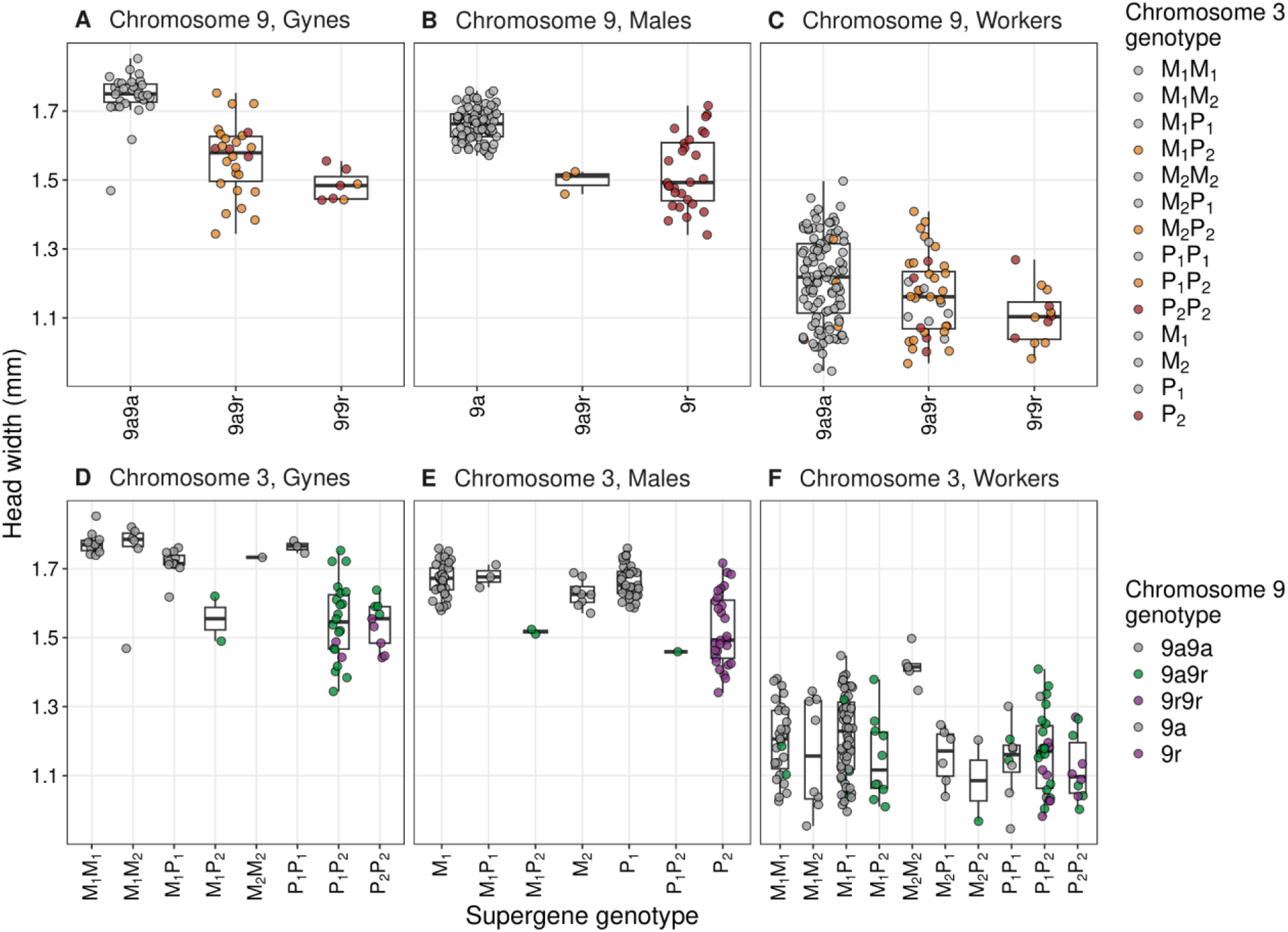
The 9r and P_2_ haplotypes are associated with smaller heads. (A-C) 9r is associated with reduced head width in males and gynes but not workers. (D-F) P_2_ is associated with reduced head width in gynes and males. P_1_ is not associated with head size. Each point represents one individual. Colors show the genotype of each individual. Alt Text: Boxplots showing head width vs genotype. Two plots each for males, gynes and workers; one for chromosome 3 and one for chromosome 9.

### Size reduction underlies association between supergenes

Similar to *F. cinerea*, microgynes of *F. lemani* are restricted to polygyne colonies due to strong linkage disequilibrium between the 9r haplotype on chromosome 9 and the P_2_ haplotype on chromosome 3. This follows the broader trend in ants of the association between colony social form, queen morphology, founding strategy, and other traits known as polygyny syndrome (Keller, 1993). Large queens generally possess the reserves needed for independent founding, whereas smaller newly mated queens commonly join established colonies or found them with the help of workers (Keller & Passera, 1989; Stille, 1996). Depending upon the social context, different queen sizes may be advantageous (Keller & Passera, 1989; Stille, 1996).

Independently evolved social supergenes have repeatedly been shown to link queen number and other traits associated with polygyny syndrome (Kay *et al*., 2022; Chapuisat, 2023). In *S. invicta*, queen weight, dispersal and colony founding strategy are linked to a marker on the social supergene (DeHeer *et al*., 1999; Wang *et al*., 2013). The social supergene in *Myrmica ruginodis* also underlies queen size (Sigeman *et al*., 2025b) and the social supergene in *Myrmecina graminicola* also underlies wing presence (Mona *et al*., 2025). In *F. selysi*, body and wing size are associated with the social supergene (De Gasperin *et al*., 2024). In *F. cinerea*, chromosomes 3 and 9 jointly underlie body length, thorax volume, wing area, mating patterns, and colony founding strategy (Scarparo *et al*., 2026a). Another ant species, *Temnothorax rugatulus,* also has separate supergenes controlling social form and queen size (Darras *et al*., 2026). The repeated association of social form and queen morphology via supergenes reinforces the idea that socially antagonistic selection may be operating to link these traits (i.e., Scarparo *et al*. 2023). This mirrors the classical model of sex chromosome evolution where sex chromosomes expand when selection favors suppressed recombination between sexually antagonistic traits and a sex determining locus (Turner, 1967; Rice, 1987; Charlesworth *et al*., 2005; Abbott *et al*., 2017; Kratochvíl *et al*., 2021).

We found some mismatched genotypes (9r without P_2_ or vice versa) in *F. lemani* workers but none among sexuals. We found the mismatch rate of *F. lemani* workers to be higher than in *F. cinerea* (23.5% vs 11.8%). In *F. cinerea*, epistatic interactions rather than physical linkage underlie the linkage disequilibrium of P_2_ and 9r (Scarparo *et al*., 2026b). Weaker epistatic interactions in *F. lemani* could cause the weaker linkage disequilibrium we observed. The magnitude of size reduction in *F. lemani* is weaker than in *F. cinerea* (14.5% vs 20%) (Scarparo *et al*., 2023), which may reduce the selective pressure for linkage disequilibrium in *F. lemani*.

Given that 9r is associated with reduced queen and male size, but not worker size, the consequences of genetic mismatches in workers may be lower. However, although non-significant, 9a9r and 9r9r workers were on average 3.3% and 7.4% smaller than 9a9a workers respectively. The lack of mismatches in sexuals could suggest high fitness consequences for sexuals, and/or a developmental bias for mismatched individuals to become workers rather than gynes.

Unlike in *F. cinerea*, we did not find P_1_ to be significantly associated with head width in *F. lemani*. In *F. cinerea*, 9a9a gynes with at least one P_1_ haplotype are 4.9% smaller than gynes without a P_1_, and P_1_ males are 2.6% larger than M_A_ males (Scarparo *et al*., 2023). In *F. lemani*, we found no significant association between P_1_ and head width. P_1_ males have the same average size as M_1_ males (Table S6). P_1_ appears to have lost its association with body size gynes as well, although we may lack the power to detect the effect size reported in *F. cinerea* because our sample size was small for P_1_P_1_ (n=3), M_1_P_1_ (n=11), and M_1_M_1_ (n=10) gynes (Table S1).

### Expanded chromosome 3 haplotype diversity

We identified four haplotypes for the social supergene in *F. lemani* (M_1_, M_2_, P_1_ and P_2_) and found all ten possible genotype combinations (Fig. 1). More than two haplotypes can occur in one system through processes such as successive chromosomal rearrangements, as in *Heliconius numata* (Joron *et al*., 2011), or via recombination between existing haplotypes, as in the butterfly *D. chrysippus* (De-Kayne *et al*., 2025) or the bird *Philomachus pugnax* (Lamichhaney *et al*., 2016). In *Formica*, recombination between the M and P haplotype has likely given rise to the M_D_ haplotype in at least two lineages (Lagunas-Robles *et al*., 2026).The M_D_ haplotype is associated with colony-level production of female sexuals in *F. cinerea* (Scarparo *et al*., 2023) and other species (Lagunas-Robles *et al*., 2021, 2026).

In *F. lemani*, M_2_ does not match the expected patterns of differentiation and ancestry we would expect from an M_D_ haplotype. M_2_ is differentiated from M_1_ across most of the supergene (2 Mbp -12.5 Mbp) while M_D_ is only differentiated from M_A_ across the first half of the supergene (∼2.67 Mbp -7.5 Mbp) (Scarparo *et al*., 2023; Lagunas-Robles *et al*., 2026). In *F. cinerea*, the M_D_ likely originated from recombination between M_A_ and P_1_ (Lagunas-Robles *et al*., 2026). The windowed PCA did not identify an interval in which M_2_ was consistently P-like, and topology weighting showed that P_1_ grouped with M_2_ and M_1_ at nearly equal frequencies (Fig. S6). While the linkage map identified a region of low recombination which aligns with the M_D_ (∼2.67-7.5 Mb) (Fig. S7; Scarparo *et al*., 2023; Lagunas-Robles *et al*., 2026), the offspring in this cross were produced by an M_2_M_2_ queen, indicating that there is low recombination *within* this region of the M_2_ haplotype, rather than suppressed recombination between alternate haplotypes. We therefore consider M_2_ a distinct M haplotype with a different evolutionary origin than M_D_.

## Conclusion

In *Formica lemani*, the supergene on chromosome 3 underlies colony queen number, but the prevalence of the P_1_ haplotype in single-queen colonies represents a significant departure from associations in other *Formica* species. This suggests that *F. lemani’s* P_1_ has acquired novel combinations of SNPs which allow compatibility with monogyny. Our analyses identified a few previously described candidate genes that could contribute to the queen number polymorphism, and detected several new ones. This suggests that several different genes may be important for queen number across the *Formica* genus. We also found evidence of a miniaturizing supergene that is in linkage disequilibrium with the social supergene, as previously observed in *F. cinerea*. The magnitude of size reduction and the magnitude of linkage disequilibrium is, however, weaker in *F. lemani*. Together, these results show that ancient supergenes are not static genetic units. Rare exchange can alter the phenotype associated with an established haplotype and selection can couple supergenes on different chromosomes when their effects are favored in the same social environment. *F. lemani* therefore illustrates how supergene systems can continue to evolve through both sequence exchange within rearrangements and functional association among rearrangements.

## Acknowledgments

We thank the members of the Brelsford and Purcell labs including Alyssa Canova, Evan Shimota, Rachel Zeng, Daniel Pierce, Marie Palanchon, Alan Wu, and Carolina Gonzalez for help in the lab. We also thank Marie Palanchon, Thomas Nonacs, and Samantha Sudoko for comments on manuscript drafts.

## Data Availability

The raw reads for the RADseq dataset will be available at NCBI under BioProject PRJNA1512539 and for the whole genome resequencing dataset will be available under BioProject PRJNA1537357 upon publication. Head width measurements can be found in table S1.

## Author Contributions

Zul Alam (Conceptualization, Formal Analyses, Investigation, Visualization, Writing-original draft, and Writing-review & editing). Giulia Scarparo (Conceptualization, Formal Analyses , Investigation, Resources, Supervision, Visualization , and Writing-review & editing). German Lagunas-Robles (Formal analysis, Investigation, and Writing-review & editing). Daniela Zarate (Formal analysis and Writing-review & editing). Jessica Purcell (Conceptualization, Formal Analyses, Funding Acquisition, Investigation, Supervision, Resources, and Writing-review & editing). Alan Brelsford (Conceptualization, Formal Analyses, Funding Acquisition, Investigation, Supervision, Resources, and Writing-review & editing).

## Literature Cited

1. Abbott, J.K., Nordén, A.K. & Hansson, B. 2017. Sex chromosome evolution: historical insights and future perspectives. Proc. R. Soc. B Biol. Sci. 284: 20162806.

2. Basler, K. & Hafen, E. 1988. Control of photoreceptor cell fate by the sevenless protein requires a functional tyrosine kinase domain. Cell 54: 299–311.

3. Berdan, E.L., Blanckaert, A., Butlin, R.K. & Bank, C. 2021. Deleterious mutation accumulation and the long-term fate of chromosomal inversions. PLOS Genet. 17: e1009411.

4. Blumer, L.M., Good, J.M. & Durbin, R. 2025. WinPCA: a package for windowed principal component analysis. Bioinformatics 41: btaf529.

5. Borowiec, M.L., Cover, S.P. & Rabeling, C. 2021. The evolution of social parasitism in *Formica* ants revealed by a global phylogeny. Proc. Natl. Acad. Sci. 118: e2026029118.

6. Boulain, H., Deshmukh, R., Avril, A., Blacher, P., Bourquin, M., Yek, S.H., et al. 2026. Ancient but dynamic: structural expansion, massive gene duplication, and transposable element colonization in a supergene controlling ant social organization. Mol. Biol. Evol. 43: msag127.

7. Brelsford, A., Dufresnes, C. & Perrin, N. 2016. High-density sex-specific linkage maps of a European tree frog (*Hyla arborea*) identify the sex chromosome without information on offspring sex. Heredity 116: 177–181.

8. Brelsford, A., Purcell, J., Avril, A., Tran Van, P., Zhang, J., Brütsch, T., et al. 2020. An ancient and eroded social supergene is widespread across *Formica* ants. Curr. Biol. 30: 304–311.e4.

9. Catchen, J., Hohenlohe, P.A., Bassham, S., Amores, A. & Cresko, W.A. 2013. Stacks: an analysis tool set for population genomics. Mol. Ecol. 22: 3124–3140.

10. Catchen, J.M., Amores, A., Hohenlohe, P., Cresko, W. & Postlethwait, J.H. 2011. Stacks: building and genotyping loci de novo from short-read sequences. G3 Genes Genomes Genet. 1: 171–182.

11. Chapuisat, M. 2023. Supergenes as drivers of ant evolution. Myrmecol. News 33: 1–18.

12. Charlesworth, B. 2023. The effects of inversion polymorphisms on patterns of neutral genetic diversity. Genetics 224: iyad116.

13. Charlesworth, D., Charlesworth, B. & Marais, G. 2005. Steps in the evolution of heteromorphic sex chromosomes. Heredity 95: 118–128.

14. Cingolani, P., Platts, A., Wang, L.L., Coon, M., Nguyen, T., Wang, L., et al. 2012. A program for annotating and predicting the effects of single nucleotide polymorphisms, SnpEff: SNPs in the genome of *Drosophila melanogaster* strain w^1118^ ; iso-2; iso-3. Fly (Austin) 6: 80–92.

15. Danecek, P., Auton, A., Abecasis, G., Albers, C.A., Banks, E., DePristo, M.A., et al. 2011. The variant call format and VCFtools. Bioinformatics 27: 2156–2158.

16. Danecek, P., Bonfield, J.K., Liddle, J., Marshall, J., Ohan, V., Pollard, M.O., et al. 2021. Twelve years of SAMtools and BCFtools. GigaScience 10: giab008.

17. Darras, H., Yu, C., Kever, M., Stoldt, M., Feldmeyer, B. & Foitzik, S. 2026. Sequential origin of an alternative colony founding strategy driven by two supergenes in *Temnothorax* ants. Mol. Biol. Evol. 43: msag115.

18. De Gasperin, O., Blacher, P., Sarton-Lohéac, S., Grasso, G., Corliss, M.K., Nicole, S., et al. 2024. A supergene controlling social structure in Alpine ants also affects the dispersal ability and fecundity of each sex. Proc. R. Soc. B Biol. Sci. 291: 20240494.

19. DeHeer, C.J., Goodisman, M.A.D. & Ross, K.G. 1999. Queen dispersal strategies in the multiple-queen form of the fire ant *Solenopsis invicta*. Am. Nat. 153: 660–675.

20. De-Kayne, R., Gordon, I.J., Terblanche, R.F., Collins, S., Omufwoko, K.S., Martins, D.J., et al. 2025. Incomplete recombination suppression fuels extensive haplotype diversity in a butterfly colour pattern supergene. PLOS Biol. 23: e3003043.

21. Delaney, S.J., Hayward, D.C., Barleben, F., Fischbach, K.F. & Miklos, G.L. 1991. Molecular cloning and analysis of small optic lobes, a structural brain gene of *Drosophila melanogaster*. Proc. Natl. Acad. Sci. 88: 7214–7218.

22. Dialynas, G., Speese, S., Budnik, V., Geyer, P.K. & Wallrath, L.L. 2010. The role of *Drosophila* Lamin C in muscle function and gene expression. Development 137: 3067–3077.

23. Dixon, G., Kitano, J. & Kirkpatrick, M. 2019. The origin of a new sex chromosome by introgression between two stickleback fishes. Mol. Biol. Evol. 36: 28–38.

24. Eyre-Walker, A. & Keightley, P.D. 2007. The distribution of fitness effects of new mutations. Nat. Rev. Genet. 8: 610–618.

25. Fox, J. & Weisberg, S. 2019. An R companion to applied regression, 3rd ed. Sage, Thousand Oaks CA.

26. Gadau, J. 2009. Phase-unknown linkage mapping in ants. Cold Spring Harb. Protoc. 2009: pdb.prot5251.

27. Guerrero, R.F., Rousset, F. & Kirkpatrick, M. 2012. Coalescent patterns for chromosomal inversions in divergent populations. Philos. Trans. R. Soc. B Biol. Sci. 367: 430–438.

28. Gutiérrez-Valencia, J., Fracassetti, M., Berdan, E.L., Bunikis, I., Soler, L., Dainat, J., et al. 2022. Genomic analyses of the *Linum* distyly supergene reveal convergent evolution at the molecular level. Curr. Biol. 32: 4360–4371.e6.

29. Hartmann, C., Landgraf, M., Bate, M. & Jäckle, H. 1997. Krüppel target gene knockout participates in the proper innervation of a specific set of *Drosophila* larval muscles. EMBO J. 16: 5299–5309.

30. Helleu, Q., Roux, C., Ross, K.G. & Keller, L. 2022. Radiation and hybridization underpin the spread of the fire ant social supergene. Proc. Natl. Acad. Sci. 119: e2201040119.

31. Jay, P., Chouteau, M., Whibley, A., Bastide, H., Parrinello, H., Llaurens, V., et al. 2021. Mutation load at a mimicry supergene sheds new light on the evolution of inversion polymorphisms. Nat. Genet. 53: 288–293.

32. Jones, B.M., Waugh, A.H., Catto, M.A., Kay, S., Glastad, K.M., Goodisman, M.A.D., et al. 2025. The fire ant social chromosome exerts a major influence on genome regulation. Mol. Biol. Evol. 42: msaf112.

33. Joron, M., Frezal, L., Jones, R.T., Chamberlain, N.L., Lee, S.F., Haag, C.R., et al. 2011. Chromosomal rearrangements maintain a polymorphic supergene controlling butterfly mimicry. Nature 477: 203–206.

34. Kay, T., Helleu, Q. & Keller, L. 2022. Iterative evolution of supergene-based social polymorphism in ants. Philos. Trans. R. Soc. B Biol. Sci. 377: 20210196.

35. Keller, L. 1993. The assessment of reproductive success of queens in ants and other social insects. Oikos 67: 177–180.

36. Keller, L. & Passera, L. 1989. Size and fat content of gynes in relation to the mode of colony founding in ants (Hymenoptera; Formicidae). Oecologia 80: 236–240.

37. Keller, L. & Ross, K.G. 1998. Selfish genes: a green beard in the red fire ant. Nature 394: 573– 575.

38. Kimura, M. 1968. Evolutionary rate at the molecular level. Nature 217: 624–626.

39. Kramer, J.M., Kochinke, K., Oortveld, M.A.W., Marks, H., Kramer, D., Jong, E.K. de, et al. 2011. Epigenetic regulation of learning and memory by *Drosophila* EHMT/G9a. PLOS Biol. 9: e1000569.

40. Kratochvíl, L., Stöck, M., Rovatsos, M., Bullejos, M., Herpin, A., Jeffries, D.L., et al. 2021. Expanding the classical paradigm: what we have learnt from vertebrates about sex chromosome evolution. Philos. Trans. R. Soc. B Biol. Sci. 376: 20200097.

41. Kulmuni, J., Seifert, B. & Pamilo, P. 2010. Segregation distortion causes large-scale differences between male and female genomes in hybrid ants. Proc. Natl. Acad. Sci. 107: 7371– 7376.

42. Kunte, K., Zhang, W., Tenger-Trolander, A., Palmer, D.H., Martin, A., Reed, R.D., et al. 2014. doublesex is a mimicry supergene. Nature 507: 229–232.

43. Lagunas-Robles, G., Alam, Z. & Brelsford, A. 2025. Unexpected absence of a multiple-queen supergene haplotype from supercolonial populations of Formica ants. J. Evol. Biol. 38: 543–553.

44. Lagunas-Robles, G., Purcell, J. & Brelsford, A. 2021. Linked supergenes underlie split sex ratio and social organization in an ant. Proc. Natl. Acad. Sci. 118: e2101427118.

45. Lagunas-Robles, G., Purcell, J., Shimota, E. & Brelsford, A. 2026. Multiple origins of a sex ratio supergene in *Formica* ants. Mol. Biol. Evol. 43: msag135.

46. Lajmi, A., Cohen, P., Lee, C.-C., Frenkel, Z., Pellen, Y. & Privman, E. 2026. Repeated evolution of supergenes on an ancient social chromosome. Curr. Biol. 36: 1233–1246.e4.

47. Lamichhaney, S., Fan, G., Widemo, F., Gunnarsson, U., Thalmann, D.S., Hoeppner, M.P., et al. 2016. Structural genomic changes underlie alternative reproductive strategies in the ruff (*Philomachus pugnax*). Nat. Genet. 48: 84–88.

48. Lenth, R.V. & Piaskowski, J. 2025. emmeans: estimated marginal means, aka least-squares means. R package version 2.0.0. https://CRAN.R-project.org/package=emmeans.

49. Li, H. 2011. A statistical framework for SNP calling, mutation discovery, association mapping and population genetical parameter estimation from sequencing data. Bioinformatics 27: 2987–2993.

50. Li, J., Cocker, J.M., Wright, J., Webster, M.A., McMullan, M., Dyer, S., et al. 2016. Genetic architecture and evolution of the S locus supergene in *Primula vulgaris*. Nat. Plants 2: 16188.

51. Martin, S.H. 2026. A model-free method for genealogical inference without phasing and its application for topology weighting. Genetics 232: iyaf181.

52. Matschiner, M., Barth, J.M.I., Tørresen, O.K., Star, B., Baalsrud, H.T., Brieuc, M.S.O., et al. 2022. Supergene origin and maintenance in Atlantic cod. *Nat*. Ecol. Evol. 6: 469–481.

53. McGuire, D., Sankovitz, M. & Purcell, J. 2022. A novel distribution of supergene genotypes is present in the socially polymorphic ant *Formica neoclara*. BMC Ecol. Evol. 22: 47.

54. Meunier, J. & Chapuisat, M. 2009. The determinants of queen size in a socially polymorphic ant. J. Evol. Biol. 22: 1906–1913.

55. Meyer, L., Barry, P., Riquet, F., Foote, A., Der Sarkissian, C., Cunha, R.L., et al. 2024. Divergence and gene flow history at two large chromosomal inversions underlying ecotype differentiation in the long-snouted seahorse. Mol. Ecol. 33: e17277.

56. Mi, H., Muruganujan, A., Huang, X., Ebert, D., Mills, C., Guo, X., et al. 2019. Protocol update for large-scale genome and gene function analysis with the PANTHER classification system (v.14.0). Nat. Protoc. 14: 703–721.

57. Mona, S., Gay, E.J., Taupenot, A., Ducancel, J., Laso-Jadart, R., Helleu, Q., et al. 2025. Genomic evidence of a complex supergene system linking dispersal to social polymorphism. Curr. Biol. 35: 6155–6162.e5.

58. Navarro, A., Barbadilla, A. & Ruiz, A. 2000. Effect of inversion polymorphism on the neutral nucleotide variability of linked chromosomal regions in *Drosophila*. Genetics 155: 685– 698.

59. Navarro, A., Betrán, E., Barbadilla, A. & Ruiz, A. 1997. Recombination and gene flux caused by gene conversion and crossing over in inversion heterokaryotypes. Genetics 146: 695– 709.

60. Nishikawa, H., Iijima, T., Kajitani, R., Yamaguchi, J., Ando, T., Suzuki, Y., et al. 2015. A genetic mechanism for female-limited Batesian mimicry in *Papilio* butterfly. Nat. Genet. 47: 405– 409.

61. Oba, Y., Sato, M., Ojika, M. & Inouye, S. 2005. Enzymatic and genetic characterization of firefly luciferase and *Drosophila* CG6178 as a fatty acyl-CoA synthetase. Biosci. Biotechnol. Biochem. 69: 819–828.

62. Pierce, D., Sun, P., Purcell, J. & Brelsford, A. 2022. A socially polymorphic *Formica* ant species exhibits a novel distribution of social supergene genotypes. J. Evol. Biol. 35: 1031–1044.

63. Pracana, R., Levantis, I., Martínez-Ruiz, C., Stolle, E., Priyam, A. & Wurm, Y. 2017. Fire ant social chromosomes: differences in number, sequence and expression of odorant binding proteins. Evol. Lett. 1: 199–210.

64. Purcell, J., Brelsford, A., Wurm, Y., Perrin, N. & Chapuisat, M. 2014. Convergent genetic architecture underlies social organization in ants. Curr. Biol. 24: 2728–2732.

65. Purcell, J., Lagunas-Robles, G., Rabeling, C., Borowiec, M.L. & Brelsford, A. 2021. The maintenance of polymorphism in an ancient social supergene. Mol. Ecol. 30: 6246– 6258.

66. Purcell, J., Scarparo, G., Lagunas-Robles, G., Sankovitz, M., West, M., Alam, Z., et al. 2025. Variation in social organization and supergene control along a latitudinal gradient. *bioRxiv*, doi: 10.1101/2025.07.20.665811.

67. Purcell, J., Zahnd, S., Athanasiades, A., Türler, R., Chapuisat, M. & Brelsford, A. 2016. Ants exhibit asymmetric hybridization in a mosaic hybrid zone. Mol. Ecol. 25: 4866–4874.

68. Purcell, S., Neale, B., Todd-Brown, K., Thomas, L., Ferreira, M.A.R., Bender, D., et al. 2007. PLINK: a tool set for whole-genome association and population-based linkage analyses. Am. J. Hum. Genet. 81: 559–575.

69. R Core Team. 2025. R: a language and environment for statistical computing. https://www.R-project.org/.

70. Rice, W.R. 1987. The accumulation of sexually antagonistic genes as a selective agent promoting the evolution of reduced recombination between primitive sex chromosomes. Evolution 41: 911–914.

71. Rohland, N. & Reich, D. 2012. Cost-effective, high-throughput DNA sequencing libraries for multiplexed target capture. Genome Res. 22: 939–946.

72. Rollmann, S.M., Mackay, T.F.C. & Anholt, R.R.H. 2005. Pinocchio, a novel protein expressed in the antenna, contributes to olfactory behavior in *Drosophila melanogaster*. J. Neurobiol. 63: 146–158.

73. Ross, K.G. & Keller, L. 1998. Genetic control of social organization in an ant. Proc. Natl. Acad. Sci. 95: 14232–14237.

74. Rosset, H. & Chapuisat, M. 2007. Alternative life-histories in a socially polymorphic ant. Evol. Ecol. 21: 577–588.

75. Satokangas, I., Nouhaud, P., Seifert, B., Punttila, P., Schultz, R., Jones, M.M., et al. 2023. Semipermeable species boundaries create opportunities for gene flow and adaptive potential. Mol. Ecol. 32: 4329–4347.

76. Scarparo, G., Brelsford, A. & Purcell, J. 2026a. Beyond queen number: two supergenes coordinate dispersal, mating, and colony founding in the ant *Formica cinerea*. *bioRxiv*, doi: 10.64898/2026.07.28.740873.

77. Scarparo, G., Lopez, A., Muro, E., Brelsford, A. & Purcell, J. 2026b. Lethal epistasis maintains strong linkage disequilibrium between unlinked supergenes. *bioRxiv*, doi: 10.64898/2026.01.08.698410.

78. Scarparo, G., Palanchon, M., Brelsford, A. & Purcell, J. 2023. Social antagonism facilitates supergene expansion in ants. Curr. Biol. 33: 5085–5095.e4.

79. Scarparo, G., West, M., Brelsford, A. & Purcell, J. 2024. Broad host use and frequent polyandry in the facultative dulotic species *Formica aserva* (Hymenoptera: Formicidae). Ann. Entomol. Soc. Am. 117: 257–269.

80. Seifert, B. 2018. The ants of Central and North Europe, 1st ed. (E. J. H. Robinson, trans). lutra, Boxberg.

81. Seifert, B., Kulmuni, J. & Pamilo, P. 2010. Independent hybrid populations of *Formica polyctena* X *rufa* wood ants (Hymenoptera: Formicidae) abound under conditions of forest fragmentation. Evol. Ecol. 24: 1219–1237.

82. Seppä, P., Helanterä, H., Chernenko, A., Trontti, K., Punttila, P. & Sundström, L. 2009. Population genetics of the black ant *Formica lemani* (Hymenoptera: Formicidae). Biol. J. Linn. Soc. 97: 247–258.

83. Sigeman, H., Satokangas, I., de Lamarre, M., Krapf, P., Nouhaud, P., Deshmukh, R., et al. 2025a. The loss of a supergene in obligately polygynous *Formica* wood ant species. Mol. Biol. Evol. 42: msaf320.

84. Sigeman, H., Seppä, P., Downing, P.A., Webster, M.T., Helanterä, H. & Viljakainen, L. 2025b. A novel supergene controls queen size and colony social organization in the ant *Myrmica ruginodis*. Mol. Biol. Evol. 42: msaf255.

85. Stille, M. 1996. Queen/worker thorax volume ratios and nest-founding strategies in ants. Oecologia 105: 87–93.

86. Sundström, L. 1995. Dispersal polymorphism and physiological condition of males and females in the ant, *Formica truncorum*. Behav. Ecol. 6: 132–139.

87. Tawdros, S., West, M. & Purcell, J. 2020. Scaling relationships in *Formica* ants with continuous worker size variation. Insectes Sociaux 67: 463–472.

88. Thomas, P.D., Ebert, D., Muruganujan, A., Mushayahama, T., Albou, L.-P. & Mi, H. 2022. PANTHER: making genome-scale phylogenetics accessible to all. Protein Sci. 31: 8–22.

89. Thompson, M.J. & Jiggins, C.D. 2014. Supergenes and their role in evolution. Heredity 113: 1– 8.

90. Turner, J.R.G. 1967. On supergenes. I. The evolution of supergenes. Am. Nat. 101: 195–221.

91. Turner, S.D. 2018. qqman: an R package for visualizing GWAS results using Q-Q and manhattan plots. J. Open Source Softw. 3: 731.

92. Vasimuddin, Md., Misra, S., Li, H. & Aluru, S. 2019. Efficient architecture-aware acceleration of BWA-MEM for multicore systems. In: 2019 IEEE International Parallel and Distributed Processing Symposium (IPDPS), pp. 314–324. IEEE, Rio de Janeiro, Brazil.

93. Vizueta, J., Xiong, Z., Ding, G., Larsen, R.S., Ran, H., Gao, Q., et al. 2025. Adaptive radiation and social evolution of the ants. Cell 188: 4828–4848.e25.

94. Wang, J. 2011. coancestry: a program for simulating, estimating and analysing relatedness and inbreeding coefficients. Mol. Ecol. Resour. 11: 141–145.

95. Wang, J., Wurm, Y., Nipitwattanaphon, M., Riba-Grognuz, O., Huang, Y.-C., Shoemaker, D., et al. 2013. A Y-like social chromosome causes alternative colony organization in fire ants. Nature 493: 664–668.

96. Wickham, H. 2016. ggplot2: elegant graphics for data analysis, 2nd ed. Springer-Verlag, New York.

97. Wilke, C. 2026. ggridges: ridgeline plots in “ggplot2.” https://CRAN.R-project.org/package=ggridges.

98. Wolf, J.I. & Seppä, P. 2016. Dispersal and mating in a size-dimorphic ant. Behav. Ecol. Sociobiol. 70: 1267–1276.

99. Wu, Y., Bhat, P.R., Close, T.J. & Lonardi, S. 2008. Efficient and accurate construction of genetic linkage maps from the minimum spanning tree of a graph. PLOS Genet. 4: e1000212.

100. Yoshida, S., Soustelle, L., Giangrande, A., Umetsu, D., Murakami, S., Yasugi, T., et al. 2005. DPP signaling controls development of the lamina glia required for retinal axon targeting in the visual system of *Drosophila*. Development 132: 4587–4598.

101. Zhang, J., Kobert, K., Flouri, T. & Stamatakis, A. 2014. PEAR: a fast and accurate Illumina Paired-End reAd mergeR. Bioinformatics 30: 614–620.

