## Supplemental Methods and Figures for "Gene flux drives functional changes in an ancient supergene in *Formica lemani*"

*P haplotype topology weighting*

We followed the topology weighting method as described in the main text, but on an independently called pseudohaploid chromosome 3 vcf consisting of 36 individuals. We included *F. lemani* $P_1$ (n = 10) and $P_2$ (n = 5) homozygotes from the whole genome sequencing data, *F. cinerea* $P_1$ (n = 10) and $P_2$ (n = 11) homozygotes, and the same *F. truncorum* male for polarization. We retained 200,017 SNPs.

*Investigation of the two M haplotypes*

We identified two distinct monogyne-associated haplotypes in our analyses. We examined whether these alternative haplotypes are homologous with the $M_A$ and $M_D$ haplotypes associated with colony sex ratio in *Formica cinerea* (Scarparo *et al.* 2023). The *F. cinerea* $M_D$ is differentiated from the $M_A$ only on the left arm of chromosome 3, and is more similar to the $P_1$ than to the $P_2$ haplotype (Lagunas-Robles *et al.* 2026). Through the following series of analyses, we concluded that the *Formica lemani* $M_2$ haplotype is not homologous with $M_D$, but is differentiated from the three other *F. lemani* haplotypes on both halves of chromosome 3.

*Differentiation between $M_1$ and $M_2$*

To examine genomic differentiation and determine the supergene boundaries between the $M_1$ and $M_2$ haplotypes in *F. lemani* we estimated $F_{ST}$ between $M_1M_1$ and $M_2M_2$ individuals from the whole genome dataset as described in the main text.

*$M_2$ Windowed PCA*

To investigate how genomic relationships change across chromosome 3, we performed a windowed PCA using the 24 *F. lemani* workers from our whole genome dataset with homozygous genotypes on chromosome 3. We independently recalled variants and refiltered using the same method as the main text and additionally retained only biallelic variants, resulting in 201,965 SNPs on Scaffold03. We then ran WinPCA (Blumer *et al.*, 2025) across chromosome 3. All four haplotypes were included in every PCA, but only $M_2$ and $P_1$ were visualized for PC1, and only $M_2$ and $P_2$ for PC2, respectively. We plotted windows containing at least 25 variants.

*$M_2$ Topology weighting*

We followed the topology weighting method as described in the main text, but on an independently called pseudohaploid chromosome 3 vcf consisting of 31 individuals. We included all *F. lemani* $M_1$ and $M_2$ homozygotes from the whole genome sequencing data, all *F. cinerea* $M_A$ and $P_1$ homozygotes, and the same *F. truncorum* male for polarization. We retained 192,202 SNPs.

*Linkage mapping*

We constructed a linkage map from $M_2M_2$ individuals from a single, monogyne colony, BGC43. We sequenced 61 workers via RadSeq following the same method as the main text. We then processed the sequencing data following the same method as the main text, but used Stacks v2.10 (Catchen *et al.*, 2013) for demultiplexing, and PEAR v0.9.11 (Zhang *et al.*, 2014) for read merging and adaptor removal, rather than the versions stated in the main text. We filtered our dataset for a minimum depth of 7 (--minDP7). We then removed 3 individuals from our dataset, one for having >25% missing data, and two for being PC1 outliers, which are potentially individuals of different ancestry. Following this, we filtered for sites with a minor allele frequency <10% (--maf 0.1), removed indels (--remove-indels), removed sites with excessive heterozygosity (over 80%), loci under 20% missing data (--max-missing 0.8), and retained only biallelic sites (min-alleles 2, --max-alleles 2). Our final dataset consisted of 58 individuals and 10,706 sites.

To construct a linkage map with MstMap (Wu *et al.*, 2008), we first converted genotype data to numerical values using vcftools (--012), then followed the protocol by Gadau (2009) which accounts for unknown phasing during linkage map construction in social insects. This protocol arbitrarily assigns a phase to every allele of each genetic marker, doubles the complete dataset, and then switches the phase of each second marker (i.e. a marker with phase 1 in the original dataset is then converted to phase 2 in the doubled dataset). In subsequent analyses, the duplicated linkage groups are discarded. We then used MstMap (Wu *et al.*, 2008) to construct the linkage map with the following parameters (population_type DH, distance_function kosambi, cut_off_p_value 0.000001, no_map_dist 30, no_map_size 1, missing_threshold 0.2, estimation_before_clustering no, objective_function ML).

**Supplemental Figures**

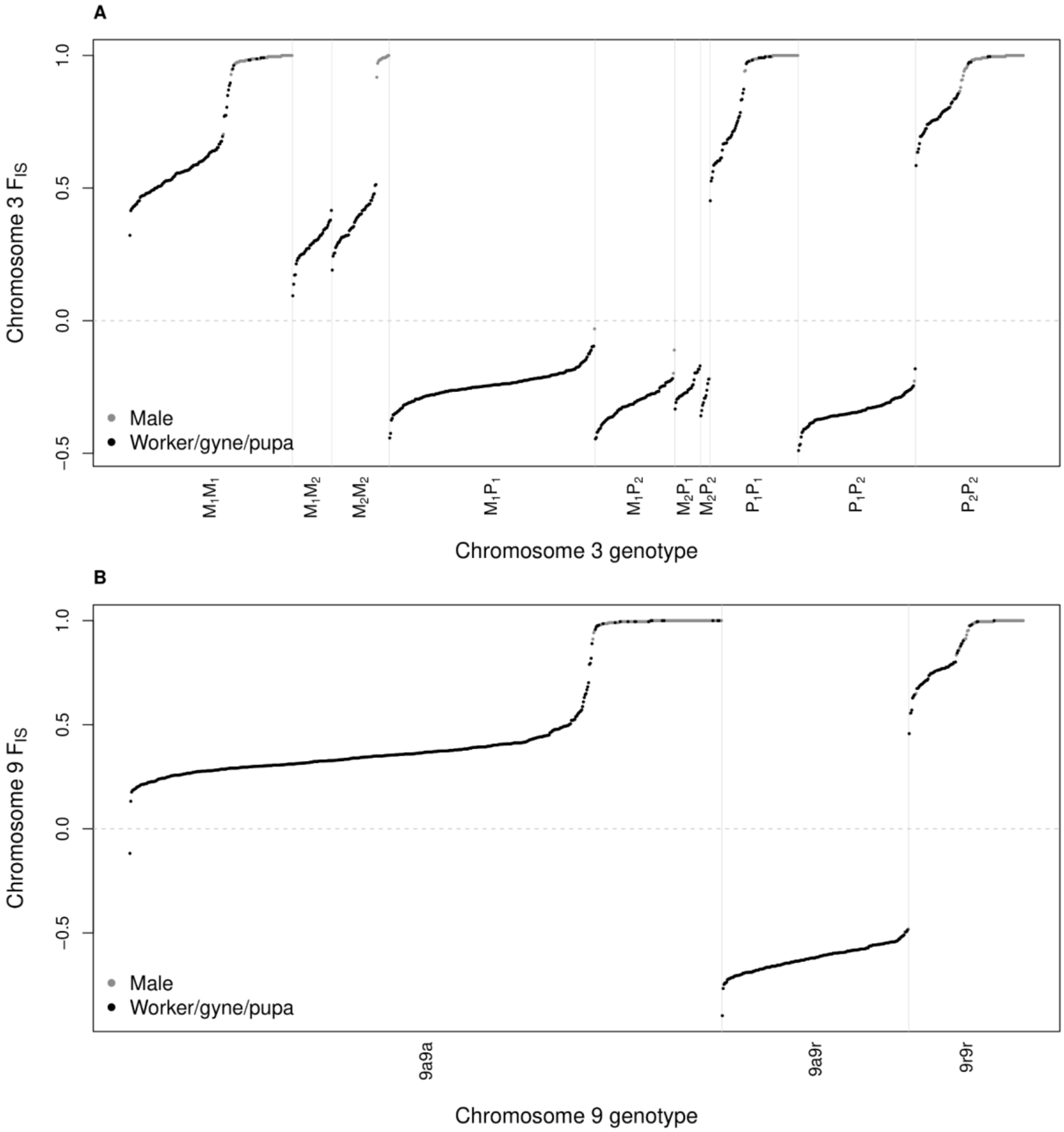


Figure S1: $F_{IS}$ calculated across chromosome 3 (A) and chromosome 9 (B) from RADseq data grouped by genotype assigned by PCA reveals that supergene heterozygotes tend to have excess SNP heterozygosity. Each dot represents one individual, black represents workers, gynes and pupae, while grey represents males. High $F_{IS}$ suggests homozygosity and low $F_{IS}$ suggests heterozygosity. High $F_{IS}$ for $M_1M_2$ is consistent with recent divergence between haplotypes. One PCA assigned 9a9a individual has a negative $F_{IS}$.

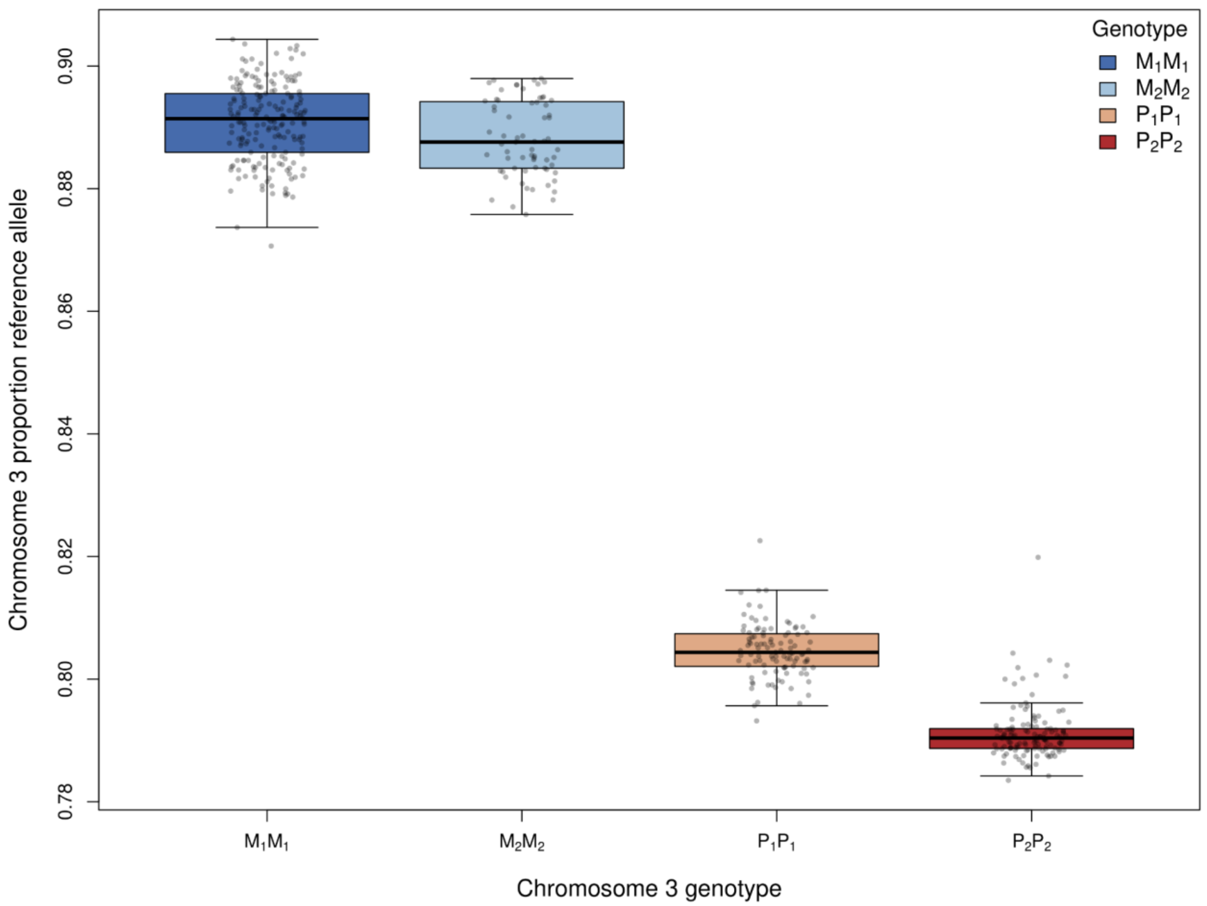


Figure S2: Proportion of reference (*F. selysi* M haplotype) alleles in *F. lemani's* chromosome 3 haplotypes show that the $M_1$ and $M_2$ are more similar to the *F. selysi* M haplotype than either P haplotype is. RADseq data was used for this figure.

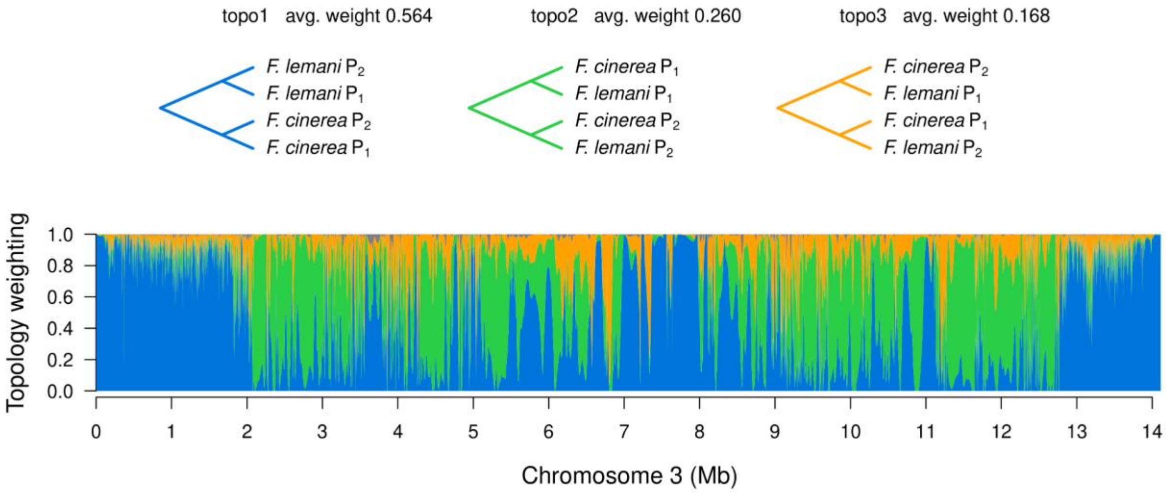


Fig S3: The *F. lemani* and *F. cinerea* $P_1$ and $P_2$ haplotypes are likely homologous. Topology weights, calculated with whole genome resequencing data, show the most common topology within the supergene region (topo2) groups the $P_1$ haplotypes together and the $P_2$ haplotypes together rather than grouping by species.

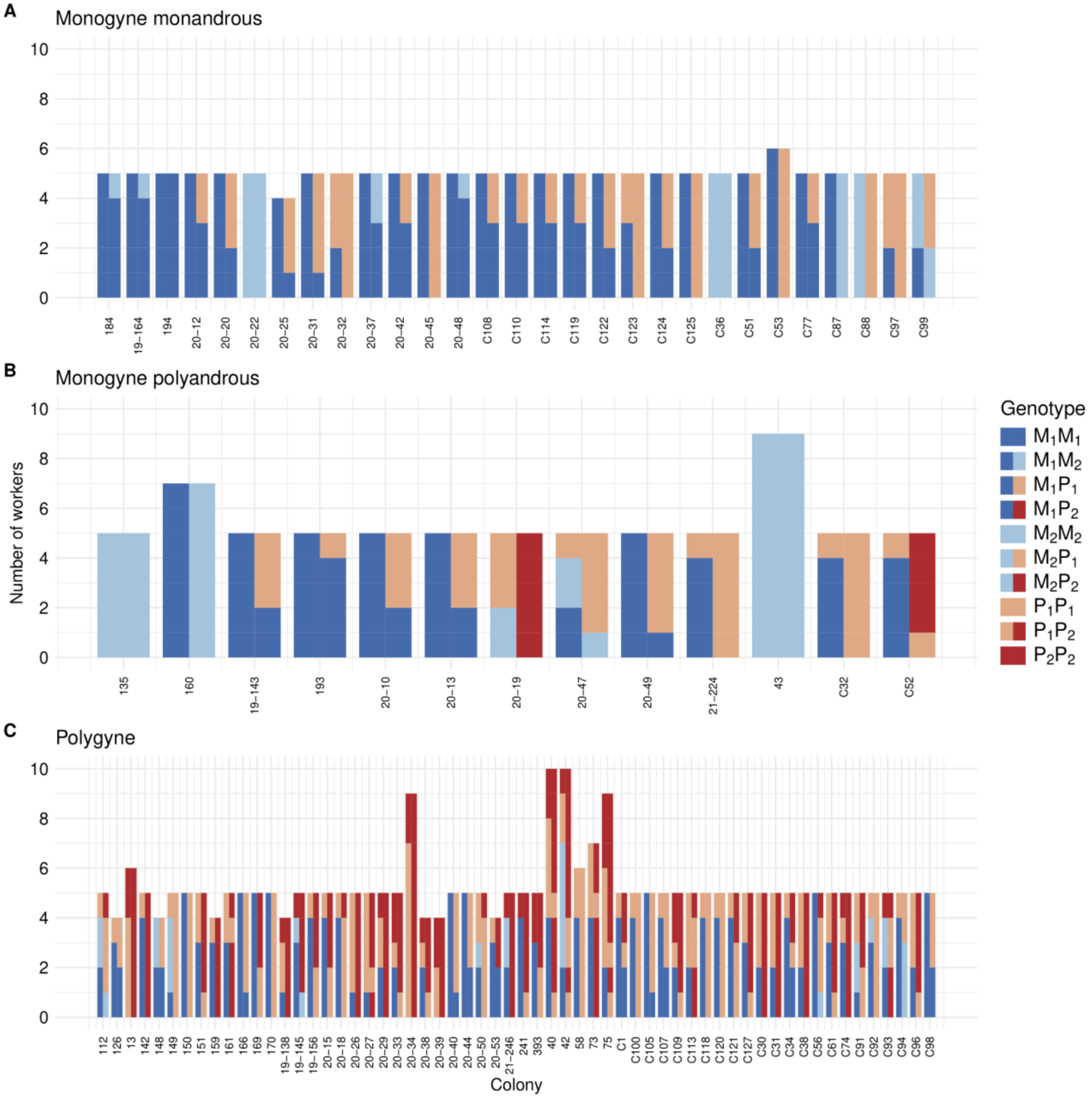


Figure S4: The haplotype counts per colony reveal that $P_2$ is nearly absent from monogyne colonies but common in polygyne colonies. The bars show the genotypes of workers from each colony. Homozygous individuals are represented by single solid colors, while heterozygous individuals have a vertical split between the colors representing their two haplotypes. (A-B) Monogyne colonies rarely have $P_2$, but $P_1$ is common. (C) $P_2$ and $P_1$ are common in polygyne colonies. All included individuals were sequenced with RADseq.

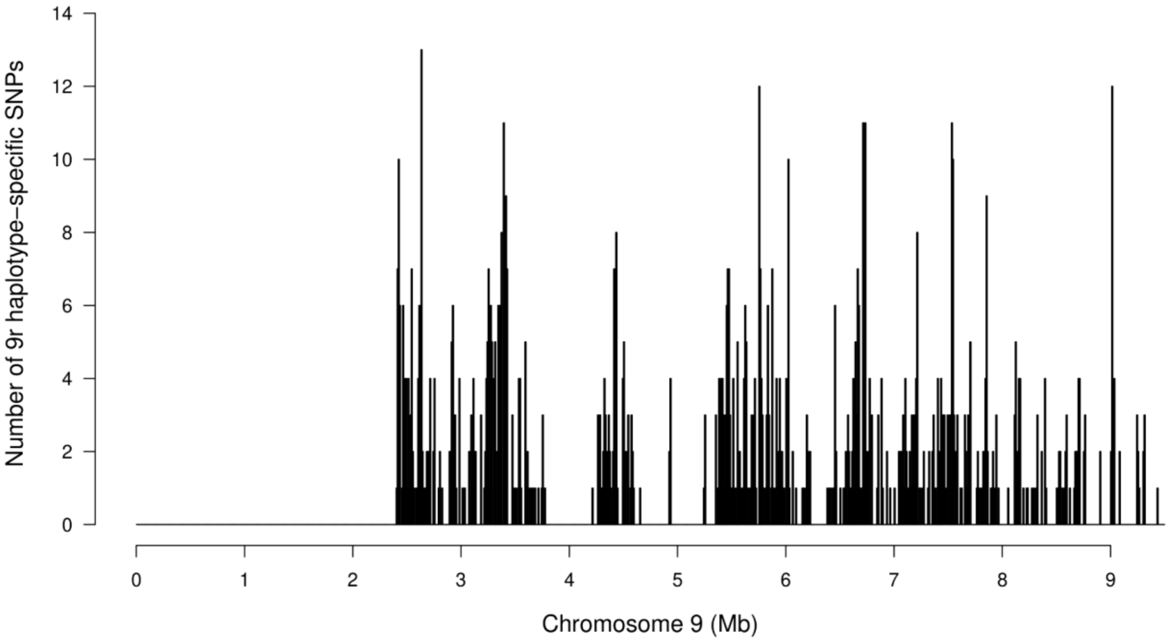


Figure S5: SNPs shared by 9r haplotypes in both *F. lemani* and *F. cinerea* are concentrated in the chromosome 9 supergene region. 925 of the shared SNPs are found in the supergene region on chromosome 9. 25 more SNPs are shared on chromosome 3, likely due to the linkage disequilibrium between 9r and $P_2$. Count per 10kb window on Chromosome 9 from whole genome resequencing data is shown.

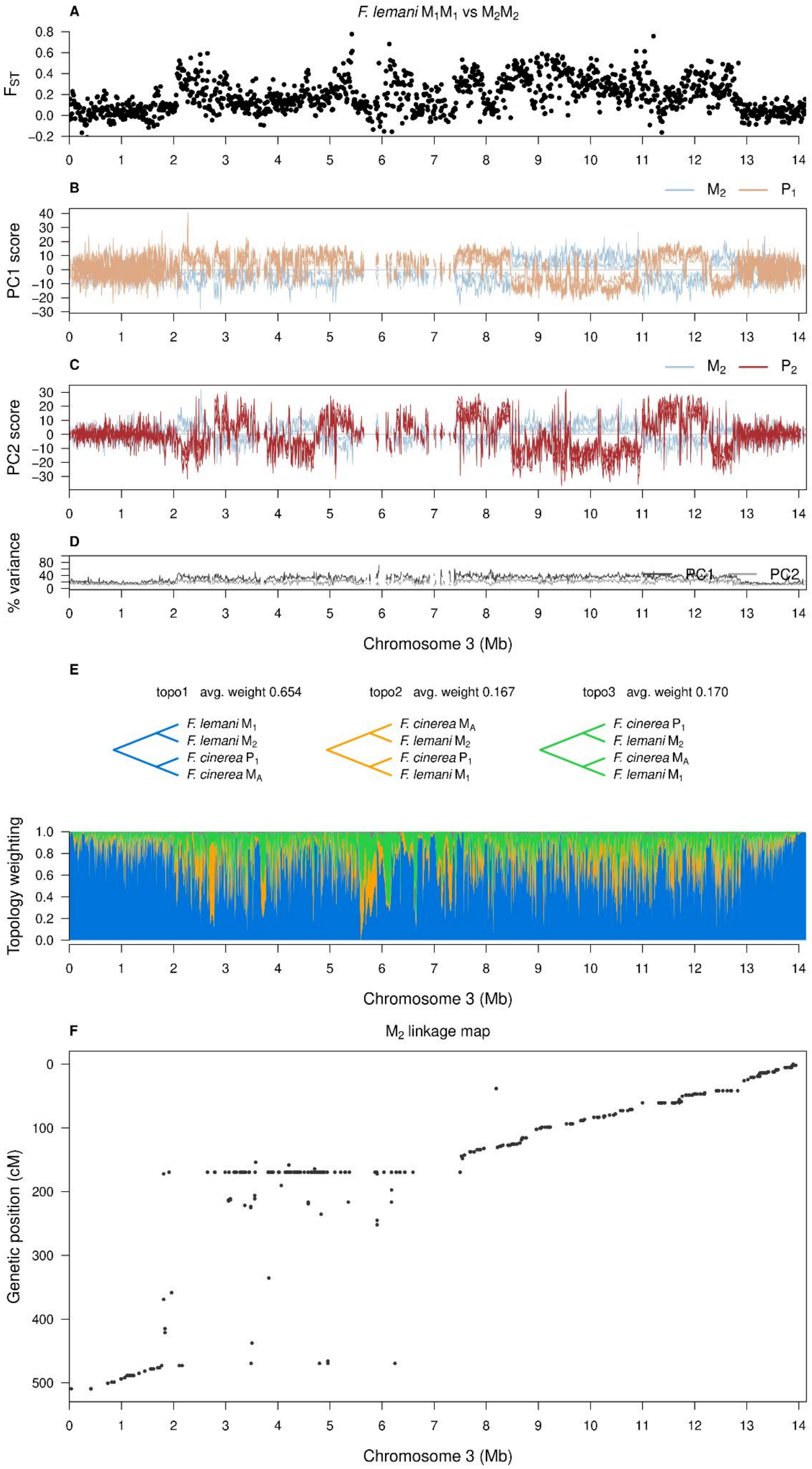

A
F. lemani M1M1 vs M2M2
FST
B
M2 P1
PC1 score
C
M2 P2
PC2 score
D
% variance
PC1 PC2
Chromosome 3 (Mb)
E
topo1 avg. weight 0.654
topo2 avg. weight 0.167
topo3 avg. weight 0.170
F. lemani M1
F. lemani M2
F. cinerea P1
F. cinerea MA
Topology weighting
Chromosome 3 (Mb)
F
M2 linkage map
Genetic position (cM)
Chromosome 3 (Mb)

Figure S6: *Formica lemani* $M_2$ is distinct from other *F. lemani* haplotypes and from *F. cinerea* $M_A$ and $M_D$. (A) $F_{ST}$ between $M_1M_1$ and $M_2M_2$ individuals shows high differentiation from ~2-12.5Mb. $F_{ST}$ calculated in 10kb windows with whole genome resequencing data. (B-C) Windowed PCA across chromosome 3 for $M_2$ and $P_1$ along PC1 (B) and $M_2$ and $P_2$ along PC2 (C) show no consistent regions where $M_2$ is similar to a P haplotype. (D) Percent variance explained by PC1 and PC2. Windowed PCAs use whole genome resequencing data. (E) Topology weights calculated with whole genome resequencing data for *F. lemani* $M_1$ and $M_2$, and *F. cinerea* $M_A$ and $P_1$ show that $M_1$ and $M_2$ group with *F. cinerea's* $P_1$ at nearly equal frequencies. (F) Linkage map of 58 $M_2M_2$ workers from a single colony show suppressed recombination from ~2.6-7.5 Mbp within the $M_2$ haplotype. Linkage map samples were sequenced via RADseq.

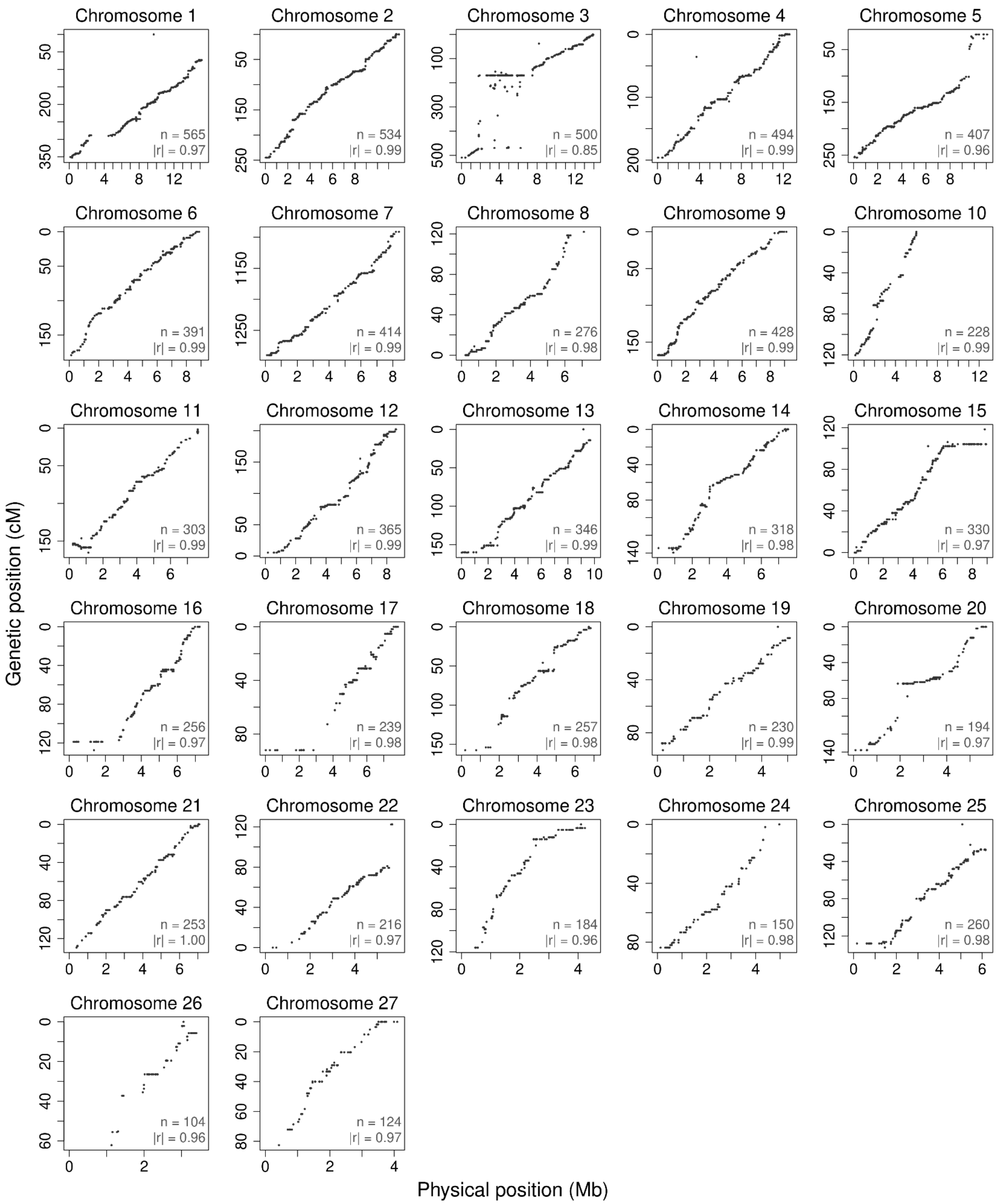


Figure S7: *F. lemani* $M_2M_2$ linkage map shows that chromosome 3 is an outlier with a large region of suppressed recombination between two copies of $M_2$ carried by the queen while most other chromosomes show a high degree of synteny between *F. lemani* and the *F. selysi* reference genome. Linkage map samples were sequenced via RADseq.
